# Human Plasma Proteomic Profile of Clonal Hematopoiesis

**DOI:** 10.1101/2023.07.25.550557

**Authors:** Zhi Yu, Amélie Vromman, Ngoc Quynh H. Nguyen, Art Schuermans, Thiago Rentz, Shamsudheen K. Vellarikkal, Md Mesbah Uddin, Abhishek Niroula, Gabriel Griffin, Michael C. Honigberg, Amy E. Lin, Christopher J. Gibson, Daniel H. Katz, Usman Tahir, Shi Fang, Sara Haidermota, Shriienidhie Ganesh, Tajmara Antoine, Joshua Weinstock, Thomas R. Austin, Vasan S. Ramachandran, Gina M. Peloso, Whitney Hornsby, Peter Ganz, JoAnn E. Manson, Bernhard Haring, Charles L. Kooperberg, Alexander P. Reiner, Joshua C. Bis, Bruce M. Psaty, Yuan-I Min, Adolfo Correa, Leslie A. Lange, Wendy S. Post, Jerome I. Rotter, Stephen S. Rich, James G. Wilson, Benjamin L. Ebert, Bing Yu, Christie M. Ballantyne, Josef Coresh, Vijay G Sankaran, Alexander G. Bick, Siddhartha Jaiswal, Robert E. Gerszten, NHLBI Trans-Omics for Precision Medicine, Peter Libby, Rajat M Gupta, Pradeep Natarajan

**Affiliations:** Broad Institute of MIT and Harvard, Cambridge, MA, USA; Cardiovascular Research Center and Center for Genomic Medicine, Massachusetts General Hospital, Boston, MA, USA; Cardiovascular Division, Brigham and Women’s Hospital Heart & Vascular Center, Boston, MA, USA; Department of Epidemiology, Human Genetics and Environmental Sciences, School of Public Health, University of Texas Health Science Center at Houston, Houston, TX, USA; Human Genetics Center, Department of Epidemiology, Human Genetics and Environmental Sciences, School of Public Health, The University of Texas Health Science Center at Houston, Houston, TX, USA; Department of Medical Oncology, Dana-Farber Cancer Institute, Boston, MA, USA; Department of Laboratory Medicine, Lund University, Lund, Sweden; Department of Pathology, Brigham and Women’s Hospital, Boston, MA, USA; Department of Pathology, Dana-Farber Cancer Institute, Boston, MA, USA; Department of Medicine, Harvard Medical School, Boston, MA, USA; Division of Cardiovascular Medicine, Beth Israel Deaconess Medical Center, Boston, MA, USA; Department of Cardiovascular Medicine, Stanford University, Stanford, CA, USA; Department of Genetics, Stanford University, Stanford, CA, USA; Department of Biomedical Engineering, Johns Hopkins University, Baltimore, MD, USA; Cardiovascular Health Research Unit, University of Washington, Seattle, WA, USA; Department of Epidemiology, University of Washington, Seattle, WA, USA; Department of Medicine, School of Medicine, Boston University, Boston, MA, USA; Framingham Heart Study, Framingham, MA, USA; Department of Biostatistics, Boston University School of Public Health, Boston, MA, USA; Division of Cardiology, Zuckerberg San Francisco General Hospital and Department of Medicine, University of California, San Francisco, CA, USA; Department of Epidemiology, Harvard T.H. Chan School of Public Health, Boston, MA, USA; Division of Preventive Medicine, Department of Medicine, Brigham and Women’s Hospital and Harvard Medical School, Boston, MA, USA; Department of Medicine III, Saarland University Hospital, Homburg, Germany; Department of Epidemiology & Population Health, Albert Einstein College of Medicine, Bronx, NY, USA; Public Health Sciences Division, Fred Hutchinson Cancer Research Center, Seattle, WA, USA; Department of Health Systems and Population Health, University of Washington, Seattle, WA, USA; Department of Medicine, University of Washington, Seattle, WA, USA; Department of Medicine, University of Mississippi Medical Center, Jackson, MS, USA; Department of Medicine, Division of Biomedical Informatics and Personalized Medicine, University of Colorado Anschutz Medical Campus, Denver, CO, USA; Division of Cardiology, Department of Medicine, Johns Hopkins University, Baltimore, MD, USA; The Institute for Translational Genomics and Population Sciences, Department of Pediatrics, The Lundquist Institute for Biomedical Innovation at Harbor-UCLA Medical Center, Torrance, CA, USA; Center for Public Health Genomics, University of Virginia, Charlottesville, VA, USA; Department of Medicine, Baylor College of Medicine, Houston, TX, USA; Department of Epidemiology, Johns Hopkins Bloomberg School of Public Health, Baltimore, MD, USA; Division of Hematology and Oncology, Boston Children’s Hospital, Harvard Medical School, Boston, MA, USA; Department of Pediatric Oncology, Dana-Farber Cancer Institute, Harvard Medical School, Boston, MA, USA; Harvard Stem Cell Institute, Cambridge, MA, USA; Department of Medicine, Vanderbilt University Medical Center, Nashville, TN, USA; Department of Pathology and Institute for Stem Cell Biology and Regenerative Medicine, Stanford University School of Medicine, Stanford, CA, USA

## Abstract

Plasma proteomic profiles associated with subclinical somatic mutations in blood cells may offer novel insights into downstream clinical consequences. Here, we explore such patterns in clonal hematopoiesis of indeterminate potential (CHIP), which is linked to several cancer and non-cancer outcomes, including coronary artery disease (CAD). Among 61,833 ancestrally diverse participants (3,881 with CHIP) from NHLBI TOPMed and UK Biobank with blood-based DNA sequencing and proteomic measurements (1,148 proteins by SomaScan in TOPMed and 2,917 proteins by Olink in UK Biobank), we identified 32 and 345 unique proteins from TOPMed and UK Biobank, respectively, associated with the most prevalent driver genes (*DNMT3A*, *TET2*, and *ASXL1*). These associations showed substantial heterogeneity by driver genes, sex, and race, and were enriched for immune response and inflammation pathways. Mendelian randomization in humans, coupled with ELISA in hematopoietic *Tet2*-/- vs wild-type mice validation, disentangled causal proteomic perturbations from *TET2* CHIP. Lastly, we identified plasma proteins shared between CHIP and CAD.

## Introduction

Clonal hematopoiesis of indeterminate potential (CHIP) is a common age-related phenomenon defined as the presence of expanded hematopoietic stem cell (HSC) clones caused by acquired leukemogenic mutations (e.g., *DNMT3A, TET2*, *ASXL1,* and *JAK2*) in persons without clinical hematologic abnormalities^1, 2^. CHIP is a pre-cancerous lesion strongly predictive of hematologic malignancy^3, 4^. In addition, CHIP predisposes an individual to other age-related human diseases, chiefly cardiovascular diseases, in both human genetic and murine experimental studies^4, 5, 6, 7, 8, 9, 10, 11, 12^.

Characterizing the consequences of CHIP mutations on the plasma proteome may facilitate an improved understanding of how CHIP influences clinical outcomes. Recent studies have associated CHIP with germline DNA variation^13, 14, 15^, bulk RNA transcript concentrations^16, 17^, and epigenomic profiles^18, 19^ for such insights. While proteins represent key downstream effector gene products, their associations with CHIP remain largely unknown. The circulating proteins are involved in numerous biological processes; surveying the proteome might offer new insights into CHIP and its mechanistic link to disease phenotypes^20^.

Leveraging paired DNA sequencing and proteomic profiling from multi-ancestry participants of four Trans-Omics for Precision Medicine (TOPMed) cohorts (N=12,911) and UK Biobank (UKB; N= 48,922), we explored the proteomic signatures of CHIP and its most common or most disease-promoting driver genes (*DNMT3A, TET2*, *ASXL1*, and *JAK2*). We prioritized potentially causal relationships with Mendelian randomization and validated this approach with ELISA studies in murine models. Lastly, we explored the functional implications of CHIP-associated proteins through pathway analyses and by examining the shared and non-shared pathways between CHIP and CAD (**Figure 1**).

**Figure 1:**
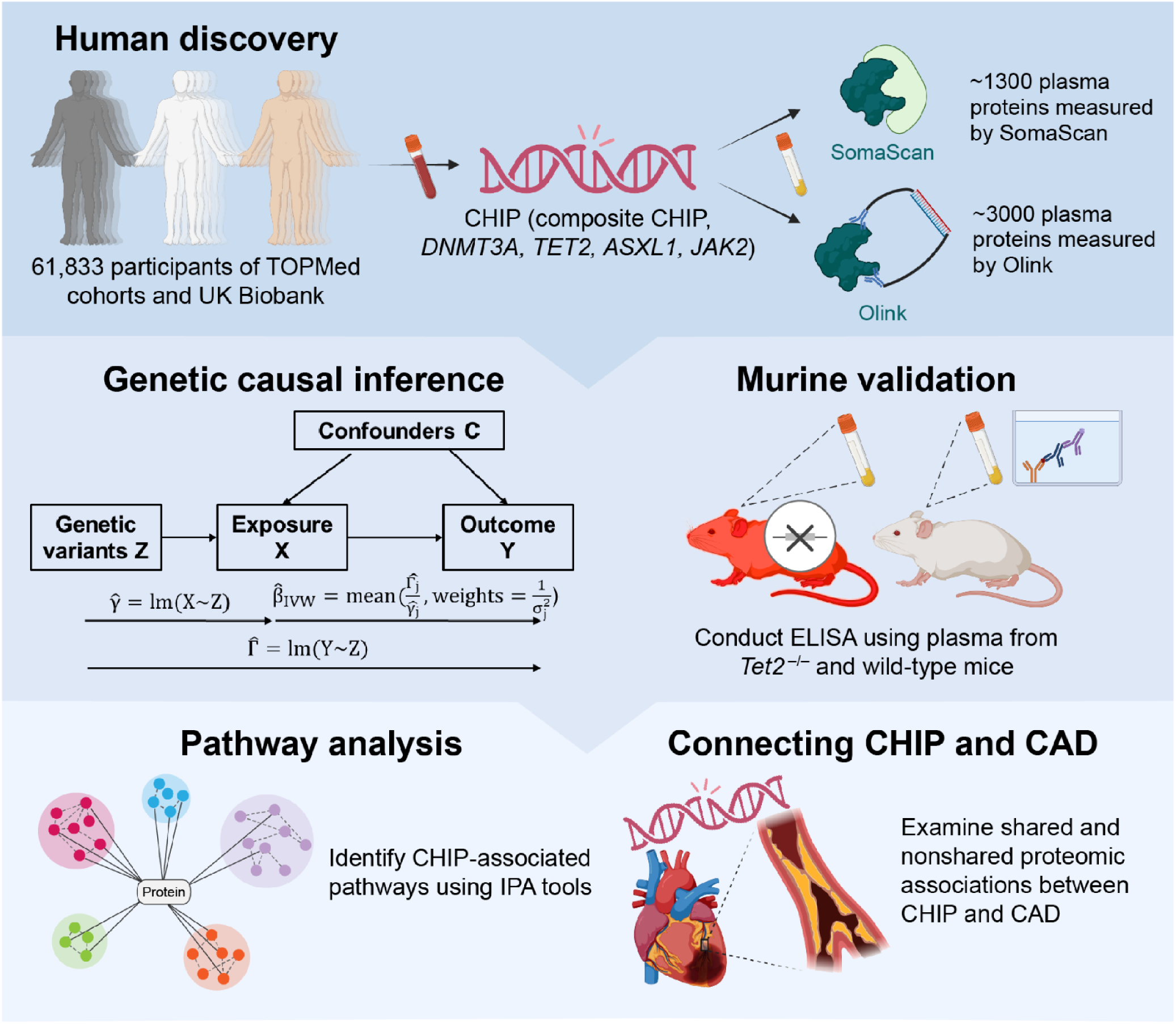
Scheme of the study design. We assessed the associations of CHIP and driver gene-specific CHIP subtypes (*DNMT3A, TET2, ASXL1*, and *JAK2*) with 1,148 circulating proteins measured by the SomaScan platform in 12,911 participants from TOPMed cohorts and 2,923 circulating proteins measured by Olink in 49,217 participants from UK Biobank. Causal relations for the associations were examined through genetic causal inference using Mendelian randomization and murine experiments contrasting plasma protein levels between *Tet2^+/+^*mice and control mice using ELISA. Pathway analyses were conducted using IPA tools. Finally, we investigated the associations between prevalent CAD and proteomics, identifying shared proteins associated with both CAD and any examined CHIP variable. CAD: Coronary artery disease. CHIP: Clonal hematopoiesis of indeterminate potential. TOPMed: ELISA: enzyme-linked immunosorbent assay. Trans-Omics for Precision Medicine. Parts of this figure have been created with BioRender.com.

## Results

### CHIP and Proteomics Characterization in Participants Across Multiple Cohorts

Our study population comprised 61,833 participants with CHIP genotyping from deep-coverage whole genome or exome sequencing of blood DNA and concurrent plasma proteomics data from four TOPMed cohorts (N=12,911), utilizing SomaScan assay for proteomics measurements^13, 21, 22^, and UK Biobank (N=48,922), whose proteomics were measured through Olink^23, 24^. The four TOPMed cohorts are the Jackson Heart Study (JHS; N=2,058)^25^, Multi-Ethnic Study of Atherosclerosis (MESA; N=976)^26^, Cardiovascular Health Study (CHS; N=1,689)^27, 28^, and Atherosclerosis Risk in Communities (ARIC; N=8,188) Study^29^.

Overviews of the study cohorts are described in detail in **Methods**. In the samples obtained from the five cohorts, 3,881 (6.0%) individuals were identified as having CHIP. Consistent with previous reports^13, 30^, CHIP was robustly associated with age. Across all cohorts, approximately 90% of individuals with CHIP driver mutations had only one identified mutation. The most commonly mutated driver genes, *DNMT3A*, *TET2*, and *ASXL1*, accounted for >75% of individuals with CHIP mutations. The variant allele fraction (VAF) distributions of mutations in each driver mutation were relatively consistent across participating cohorts (**Figure 2**, **Supplemental Tables 1-2)**.

**Figure 2.**
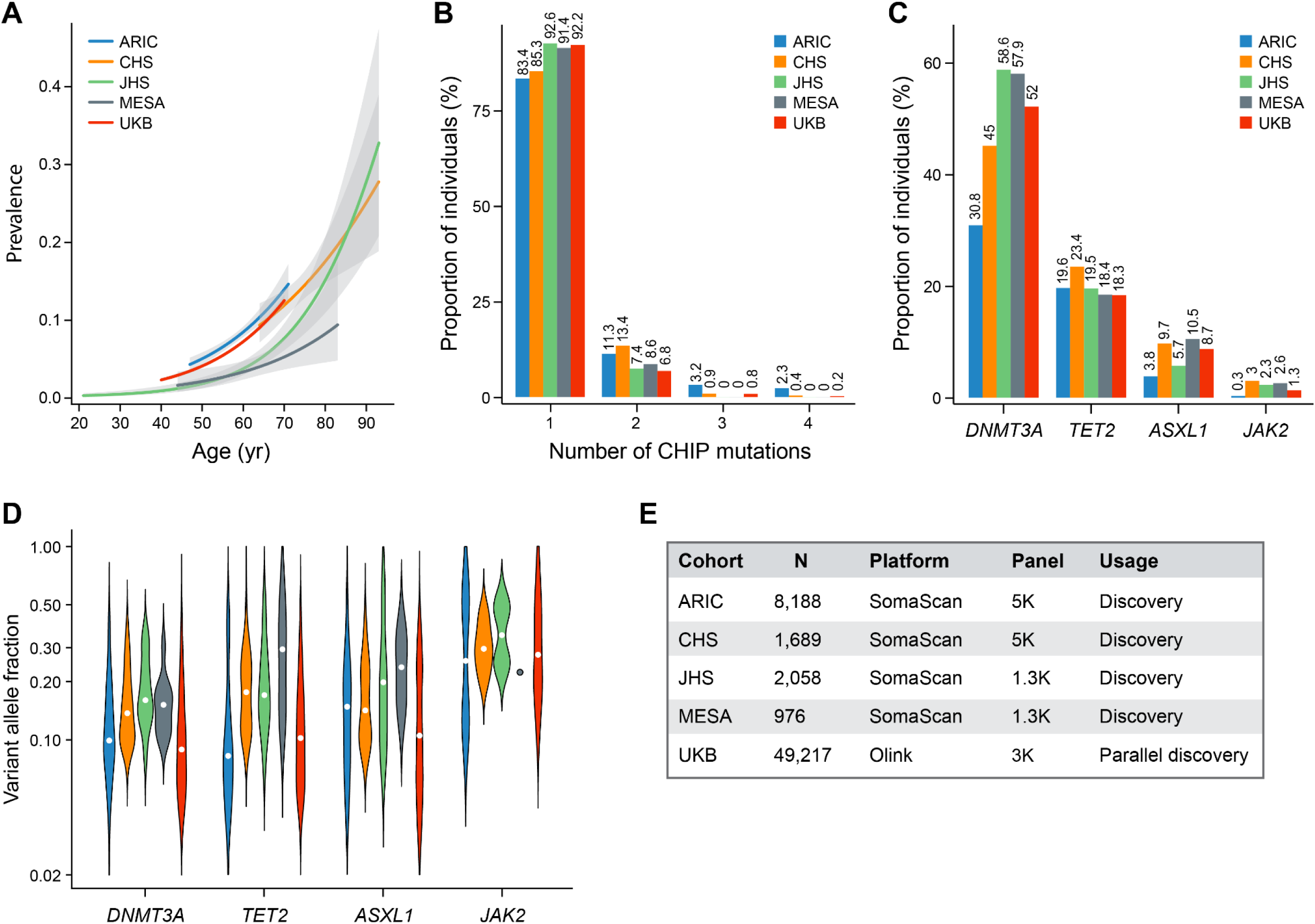
CHIP and proteomics in TOPMed cohorts and UK Biobank. A, CHIP prevalence increased with donor age at the time of blood sampling. The center line represents the general additive model spline, and the shaded region is the 95% confidence interval (N_ARIC_=8,188; N_CHS_=1,689; N_JHS_=2,058; N_MESA_=976; N_UKB_=49,217). B. More than 90% of individuals with CHIP had only one somatic CHIP driver mutation variant identified. C. Counts for four driver genes, *DNMT3A, TET2, ASXL1*, and *JAK2*, of CHIP mutations. D. CHIP clone size heterogeneity as measured by variant allele fraction by CHIP driver gene. Violin plot spanning minimum and maximum values. E. Platform and panel used for proteomics measurement by each cohort. CHIP: Clonal hematopoiesis of indeterminate potential.

### Diverse Proteomic Associations Across CHIP Driver Genes

CHIP was modeled both as a composite and separately for the most common or pathogenic drivers (*DNMT3A*, *TET2*, *ASXL1,* and *JAK2*) and defined both using the conventional thresholds for all mutations (VAF ≥2%) and for the expanded (large) clones (VAF ≥10%)^1^, resulting in ten CHIP exposure variables. As SomaScan and Olink had a relatively small overlap of proteins included in their panels with variable correlations between the overlapped proteins, we conducted separate analyses in parallel. For TOPMed cohorts that used SomaScan for proteomics measurements, the cross-sectional associations between CHIP mutations and 1,148 plasma proteins present in all cohorts were estimated within each cohort and then meta-analyzed. Consistent with prior modeling of proteomic analyses of ARIC^31^, we separated ARIC into two subpopulations: European Ancestry (EA) and African Ancestry (AA). For UK Biobank, which used Olink for proteomics measurements, we examined the cross-sectional associations between CHIP mutations and 2,917 plasma proteins in parallel. *JAK2* analyses were only conducted in cohorts with greater than 5 participants with *JAK2* mutations, which only retained CHS, ARIC EA, and UK Biobank for these analyses; thus, *JAK2* analyses were considered secondary.

Since the associations between proteins and all CHIP mutations (i.e., VAF ≥2%) are highly correlated to those with their corresponding expanded mutations (i.e., VAF ≥10%) (**Supplemental Table 3**), we retained the one with the stronger association (i.e., larger absolute Z score) to maximize power. In SomaScan-based TOPMed cohorts, this led to the identification of 35 significant CHIP variable-protein pairs (false discovery rate [FDR]<0.05, 4,592 testings), representing 32 unique proteins, independent of potential confounders described in detail in Materials and Methods (**Figure 3** and **Supplemental Table 4**). Adding *JAK2* increased the number of significant pairs to 107 (**Supplemental Figure 1** and **Supplementary Table 4**). In the Olink-based UK Biobank, 473 CHIP variable-protein pairs (345 unique proteins) passed FDR<0.05 threshold and the number increased to 861 when adding *JAK2* (**Figure 4**, **Supplemental Table 5**, **Supplemental Figure 2**).

**Figure 3.**
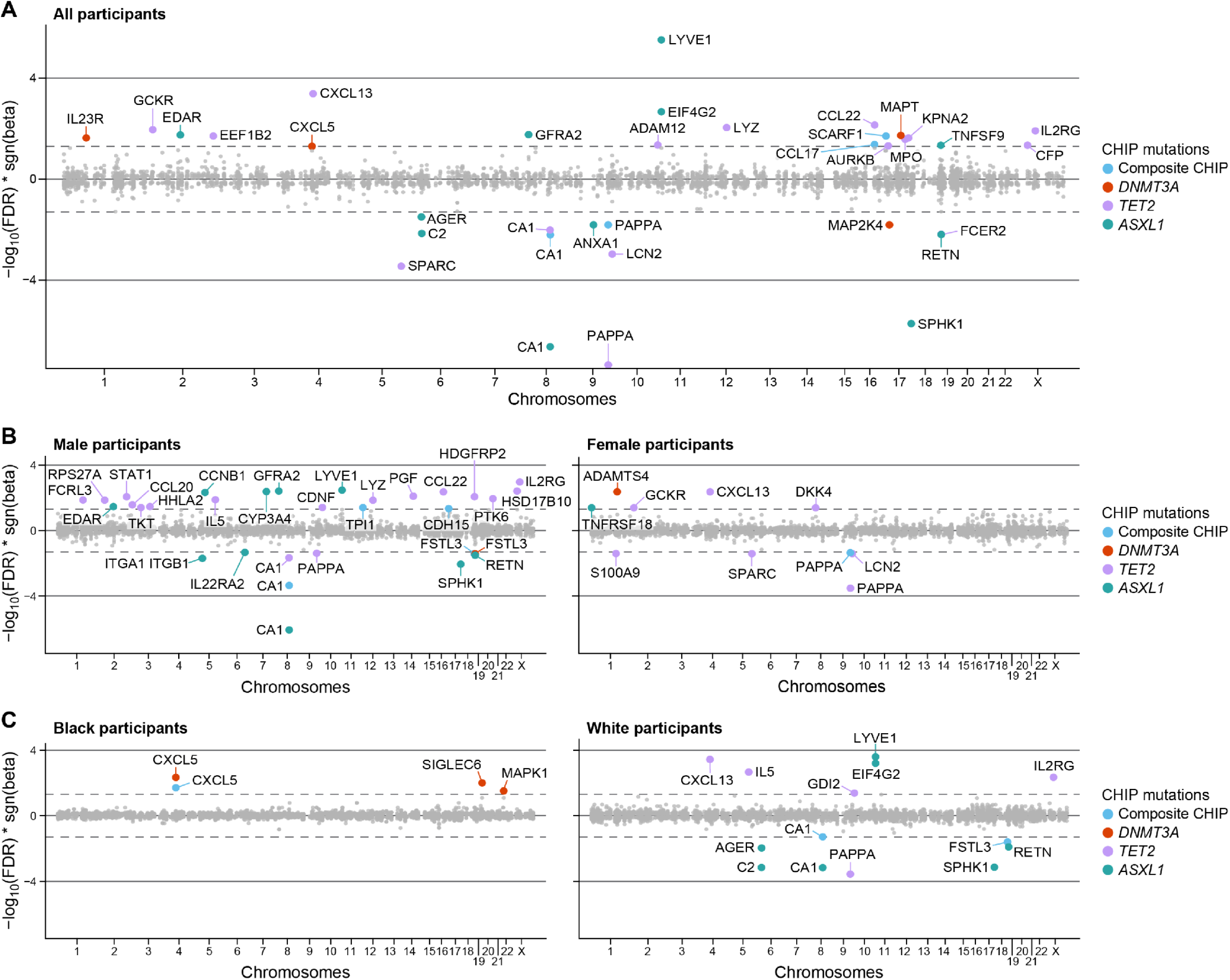
Meta-analyzed associations between CHIP mutations and circulating proteome measured by SomaScan in TOPMed cohorts. A. All participants (N=12,911). B. Male participants only (N=5,616) vs. Female participants only (N=7,295). C. Black participants only (N=4,452) vs. White participants only (N=8,076). Proteins that are associated at FDR=0.05 level (for 4,560 testings) are labeled with the corresponding SomaScan targets and colored in blue, red, green, and orange, indicating significant associations with composite CHIP, *DNMT3A*, *TET2*, and *ASXL1*, respectively. Associations were assessed through linear regression models adjusting for age at sequencing, sex (if applicable), self-reported race (if applicable), batch (if applicable), type 2 diabetes status, smoker status, first ten principal components of genetic ancestry, and PEER factors (the number of PEER factors varies by cohorts based on the sizes of study populations: 50 for JHS, MESA, and CHS; 70 for ARIC AA; 120 for ARIC EA). AA: African Ancestry; ARIC: Atherosclerosis Risk in Communities; CHIP: Clonal hematopoiesis of indeterminate potential; CHS: Cardiovascular Heart Study; EA: European Ancestry; FDR: False discovery rate; JHS: Jackson Heart Study; MESA: Multi-Ethnic Study of Atherosclerosis; PEER: Probabilistic estimation of expression residuals.

**Figure 4.**
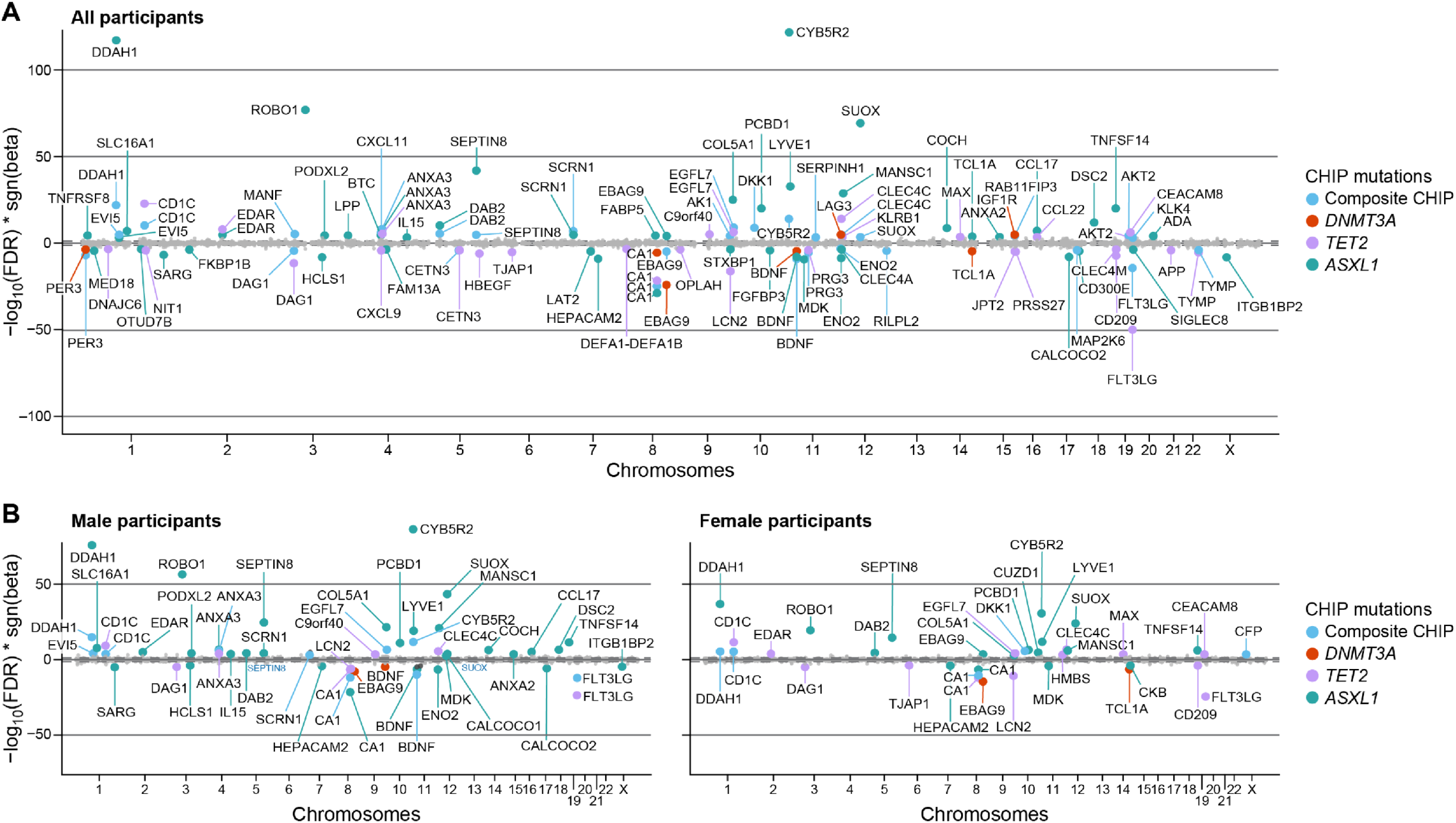
Associations between CHIP mutations and circulating proteome measured by Olink in UK Biobank. A. All participants (N=41,022). B. Male participants only (N=18,831) vs. Female participants only (N=22,191). Proteins that are associated at FDR=0.005 level (for 11,668 testings) are labeled with the corresponding Olink targets and colored in blue, red, purple, and green, indicating significant associations with composite CHIP, *DNMT3A*, *TET2*, and *ASXL1*, respectively. Associations were assessed through linear regression models adjusting for age at sequencing, sex, self-reported British White ancestry (if applicable), type 2 diabetes status, current smoker status, first ten principal components of genetic ancestry, and 150 PEER factors. CHIP: Clonal hematopoiesis of indeterminate potential; FDR: False discovery rate; PEER: Probabilistic estimation of expression residuals.

Consistent with prior work implicating heightened interleukin (IL)-1β, NOD-, LRR- and pyrin domain-containing protein 3 (NLRP3), IL-6R pathways in CHIP biology^9, 13, 16, 17, 32^, the proteins associated with examined CHIP mutations (including *JAK2*) at FDR=0.05 level were similarly enriched in these inflammatory pathways. For example, *TET2* was negatively associated with lipocalin 2 (LCN2), a secreted glycoprotein upregulated by IL-1β signaling and contributing indirectly to NLRP3 inflammasome activity in both TOPMed cohorts (SomaScan) and UK Biobank (Olink). Significant associations between CHIP variables and interleukin-related proteins involved in the pathways, such as IL-1 receptor type 1 (IL1R1) and type 2 (IL1R2), IL10, and IL-18 binding protein (IL18BP), were also observed in either TOPMed cohorts or UK Biobank. In addition, we also observed significant positive associations between CHIP variables and a number of chemokines play a role in immune cell recruitment and activation during inflammation and also contribute to the production and regulation of IL-1β and IL-6, consistently in both TOPMed cohorts and UK Biobank, such as C-C motif chemokine ligand (CCL) 17, CCL22, CCL28, C-X-C motif chemokine ligand (CXCL) 5, and CXCL11. There were additional significant associations in either study population with other chemokines, such as CXCL9, and tumor necrosis factor superfamily (TNFSF) members, such as TNFSF14 (**Figure 3**-**4**, **Supplemental Figure 1-2**, and **Supplemental Tabe 4**-**5**).

In both TOPMed cohorts and UK Biobank, mutations in individual CHIP genes exhibited distinct proteomic associations. Among the primary CHIP variables*, TET2* and *ASXL1* demonstrated a larger number of associations with plasma proteins compared with the most prevalent driver gene, *DNMT3A. TET2* was associated with 16 proteins in TOPMed cohorts and 121 proteins in UK Biobank at the FDR<0.05 threshold. *ASXL1* was associated with 11 proteins in TOPMed cohorts and 157 proteins in UK Biobank at the FDR<0.05 threshold. In contrast, *DNMT3A* was significantly associated with only four and 59 proteins in TOPMed cohorts and UK Biobank, respectively, and the associations are generally weaker than those with *TET2* and *ASXL1.* Proteins associated with composite CHIP were typically driven by individual mutant genes. Despite its infrequency and restricted sample size for secondary analysis, *JAK2* was associated with more proteins than all primary CHIP mutations examined, with 54 proteins associated in TOPMed cohorts and 315 proteins in UK Biobank. Proteins associated with *JAK2* also generally differed from other examined driver genes (**Figure 3**-**4**, **Supplemental Figure 1-2**, and **Supplemental Tabe 4**-**5**). In the TOPMed meta-analysis, some CHIP variable-protein associations have heterogeneity across cohorts, and this was mainly observed in the associations between *JAK2* and proteins.

Proteins associated with different CHIP variables are enriched with different functions. Although the proteins measured by SomaScan and Olink are different, the enriched functions showed some convergence. *TET2* was associated with proteins predominately involved in immune regulation, as well as extracellular matrix (ECM) remodeling and cell signaling. In TOPMed cohorts, for example, the top two associated proteins, pappalysin-1 (PAPPA) and secreted protein, acidic and rich in cysteine (SPARC), both participate in ECM remodeling^33, 34, 35, 36, 37, 38, 39^. Detailed protein functions are discussed in the **Supplemental Text**. For participants with *TET2* mutations, plasma PAPPA and SPARC levels were 22% and 8.2% lower, respectively, than those without *TET2* mutations (FDR = 4.6×10^-8^ and 3.8×10^-4^, respectively). We observed significant associations between *TET2* CHIP and several proteins related to immune regulation. For example, the third and fifth strongest associated proteins CXCL13 and CCL22 (both positive associations; FDR = 3.8×10^-4^ and 6.7×10^-3^, respectively) are implicated in regulating IL-1β and IL-6 levels as mentioned above, and a few other proteins are involved in innate immunity, such as lipocalin-2 (LCN2; negative association; FDR = 1.1×10^-3^; heterogeneity p value = 3.2×10^-5^ [suggesting less robust evidence]) and myeloperoxidase (MPO; positive association; FDR = 0.02) (**Figure 3** and **Supplementary Table 4**). In the UK Biobank, notably, LCN2 and SPARC, associated with TET2 in the TOPMed cohorts, show consistent associations in the UK Biobank (FDR = 9.4×10^-17^and 0.005, respectively). *TET2* is associated with fms-related tyrosine kinase 3 ligand (FLT3LG), T-cell surface glycoprotein CD1c (CD1C), C-type lectin domain family 4 member C (CLEC4C), and CD209 (FDR = 2.1×10^-50^, 1.4×10^-23^, 1.2×10^-14^, and 1.1×10^-7^, respectively). These proteins play key roles in the regulation and activation of immune responses. *TET2* is also associated with tumor necrosis factor receptor superfamily member EDAR (EDAR), epidermal growth factor-like protein 7 (EGFL7), and proheparin-binding EGF-like growth factor (HBEGF) (FDR = 1.5×10^-8^, 6.2×10^-7^, and 2.2×10^-6^, respectively), which are involved in cell growth, differentiation, and survival signaling pathways. Additionally, DAG1 and COL4A1 (FDR = 1.2×10^-14^ and 0.001, respectively) are linked to *TET2*, contributing to cell structure maintenance and extracellular matrix interactions (**Figure 4** and **Supplementary Table 5**).

In addition to immune regulation, *ASXL1*-associated proteins were enriched in metabolic regulation and cell signaling. For example, carbonic anhydrase 1 (CA1), which is crucial for metabolic processes related to pH and ion balance, is the top protein associated with *ASXL1* in TOPMed cohorts and also strongly associated with *ASXL1* in UK Biobank^40^. In both study populations, CA1 is significantly associated with other CHIP variables. In the TOPMed cohorts, CA1 levels are 15.9% (FDR = 2.4×10^-7^), 8.2% (FDR = 0.01), and 4.3% (FDR = 6.7×10^-3^) lower among participants with *ASXL1*, *TET2*, and composite CHIP, respectively, than those without those mutations. Similarly, in the UK Biobank, CA1 levels are 21.8% (FDR = 2.4×10^-29^), 12.4% (FDR = 3.2×10^-22^), 6.3% (FDR = 3.2×10^-25^), and 4.4% (FDR = 5.7×10^-6^) lower among participants with *ASXL1*, *TET2*, composite CHIP, and *DNMT3A*, respectively, than those without those mutations. The top two associated proteins with *ASXL1* in UK Biobank are cytochrome B5 reductase 2 (CYB5R2) and dimethylarginine dimethylaminohydrolase 1 (DDAH1); both are key metabolic enzymes, with the former supporting electron transport and metabolic stability and the latter regulating nitric oxide levels to promote vascular health (both positive associations; FDR = 2.1×10^-122^ for CYB5R2 and FDR = 6.8×10^-118^). Other proteins associated with *ASXL1* span metabolic regulation, immune regulation, and cell signaling. For example, metabolic protein, resistin (RETN)^41^, and immune-regulating proteins, EDAR and lymphatic vessel endothelial hyaluronic acid receptor 1 (LYVE1)^43^, are strongly associated with *ASXL1* in both TOPMed cohorts and UK Biobank with consistent directions of effects. And *ASXL1,* in TOPMed cohorts, is associated with sphingosine kinase 1 (SPHK1), a protein on signaling pathways that regulate cell growth and proliferation^42^, and, in UK Biobank, is also strongly associated with roundabout guidance receptor 1 (ROBO1), a neural cell adhesion molecule (**Figure 3-4** and **Supplementary Table 4**-**5**)^42, 43^.

In the secondary analysis, *JAK2* is associated with 54 proteins in TOPMed cohorts and 316 proteins in UK Biobank that exhibit diverse functions. The top *JAK2*-associated proteins are highly consistent between TOPMed cohorts and UK Biobank: Top associated proteins in both study populations, including P-selectin (SELP) and platelet glycoprotein 1b alpha chain (GP1BA), play crucial roles in cell adhesion and platelet function had greater concentrations among those with *JAK2*^44, 45^ similar to knock-in mice with inducible *JAK2^V617F^* ^13, 46^; additionally, individuals with *JAK2* CHIP had reduced erythropoietin (EPO) concentrations in both study populations, which has been observed among individuals with *JAK2* myeloproliferative neoplasms^47^; other top-associated proteins in both study populations are involved in bone metabolism and signaling pathway regulation (dickkopf WNT signaling pathway inhibitor 1 [DKK1]), growth and neural development (amphoterin induced gene and ORF 2 [AMIGO2]) and pleiotrophin [PTN]), and immune response (CXCL11) (**Supplemental Figure 1-2** and **Supplemental Table 4-5**)^48^. In ARIC, we additionally adjusted for platelet and white blood cell (WBC) counts in the sensitivity analysis, and the results were largely robust. The associations between *JAK2* and a few proteins directly related to platelets, such as GP1BA, were diminished but remained statistically significant (**Supplemental Table 6** and **Supplemental Figure 3**). We also investigated the relationship between the VAF of CHIP variables and proteomics in the UK Biobank, finding significant associations comparable to those observed with binary CHIP variables (**Supplemental Table 7**). For the aforementioned analysis, additionally adjusting for estimated glomerular filtration rates (eGFR) yielded consistent results (**Supplemental Table 8 and Supplemental Figure 4-5**).

### Comparative Analysis of Proteomic Associations Across Platforms

While we observed some consistent associations between results from TOPMed cohorts using SomaScan for proteomics analysis and UK Biobank, which utilized the Olink platform for proteomics measurements^21, 24^, the general agreement between the two platforms is moderate, consistent with recent report^49^. There were 493 unique proteins shared between the two platforms. Among the 2,465 CHIP variable-protein pairs being compared (493 proteins×5 CHIP variables [*DNMT3A, TET2, ASXL1, JAK2*, and composite CHIP]), 30.8% was nominally significant in at least one of the SomaScan-based and Olink-based results. Among them, 114 were nominally significant in both sets of results, with 26 of them being significant after correcting for multiple testing (FDR = 0.05) in both SomaScan-based and Olink-based results. Those pairs include the top proteins associated with *JAK2*, strong signals of *ASXL1*, composite CHIP, and *TET2* with CA1, as well as the association between *TET2* and LCN2 (**Supplemental Figure 6**). Sensitivity analysis restricted to results from EA only yielded slightly dampened but generally consistent results (**Supplemental Figure 7**). Since only 19% of overlapping proteins are highly correlated between the two platforms in prior work^50^, we can not rule out the possibility of false positives and false negatives in this cross-platform comparison.

### Sex-specific and Race-specific Differences in CHIP Variable-Protein Associations

We conducted stratified analyses by sex (both TOPMed cohorts and UK Biobank) and race (TOPMed cohorts only). While there was no difference in the prevalence of composite CHIP and each examined driver gene by sex, more proteins were associated with CHIP mutations, and the associations are generally stronger in males than in females. In TOPMed, there is a relatively small overlap between significantly associated proteins between females and males (**Supplemental Table 9-10** and **Figure 3B**). For the 40 CHIP variable-protein pairs that are only significant in male or female stratified analysis, we introduced and tested for interaction terms between the corresponding CHIP variables and sex in the combined analysis across all discovery cohorts. Of these, 15 pairs displayed statistically significant interactions at an FDR = 0.05 (**Supplemental Table 11**). The sex difference slightly dampened in UK Biobank results; although only 1/4 of proteins significant in males were also significant in females, the top associated proteins showed high consistency between males and females. Some interesting sex-specific effects were observed. For example, in females, TCL1 family AKT coactivator A (TCL1A) was significantly positively associated with *TET2* and negatively associated with *DNMT3A*, as recent GWAS discoveries. But this protein was neither associated with *TET2* nor *DNMT3A* in males (**Supplemental Table 12-13** and **Figure 4B**).

Proteins associated with CHIP mutations also differ by self-reported race. In TOPMed cohorts, among individual driver genes, only *DNMT3A* demonstrated significant proteomic associations in the Black-only analysis, with two out of three associated proteins, namely sialic acid-binding Ig-like lectin 6 (SIGLEC6) and mitogen-activated protein kinase 1 (MAK1), not observed in combined analyses. In contrast, significant associations in the White-only analyses were primarily driven by *TET2* and *ASXL1* and largely reflected findings in combined analyses. And these significant associations were not present in Black-only analyses (**Supplemental Tables 14-15** and **Figures 3C**). We tested for interaction terms between CHIP variables and race in combined analysis in ARIC for the 18 proteins that are only significant in Black or White stratified analysis. Three pairs displayed statistically significant interactions at an FDR = 0.05 (**Supplemental Table 16**). *JAK2* analyses yielded similar sex and race-specific patterns (**Supplemental Tables 9-16** and **Supplemental Figures 8-13**).

### Genetic Causal Inference for CHIP-proteomics Associations

We performed genetic causal inference for CHIP-proteomic pairs with FDR < 0.05 to disentangle the potential proteomic causes and consequences of CHIP using Mendelian randomization (MR; **Methods**). Given that only composite CHIP, *DNMT3A*, and *TET2* have adequate GWAS power, we focused on these three CHIP variables for MR analysis to minimize the influence of weak instrument bias. In the TOPMed cohorts, among the 22 pairs with valid instruments, we identified nine pairs where CHIP variables causally influence proteomic changes. The strongest genetic causal effect was composite CHIP on scavenger receptor class F member 1 (SCARF1), with composite CHIP presence leading to a 7% increase in SCARF1 levels. Other significant effects included CHIP on PAPPA and *TET2* on MPO. Additionally, we found one pair where a protein level difference influenced the development of a CHIP variable: higher lysozyme (LYZ) levels decreased the risk of developing *TET2* mutations. In the UK Biobank, we observed 121 out of 317 pairs where CHIP variables causally influenced proteomic changes. Notable effects included the causal impact of CHIP and *TET2* on decreased FLT3LG concentrations and *TET2*’s causal effect on LCN2 (**Figure 5A-B** and **Supplemental Table 17- 20**).

**Figure 5.**
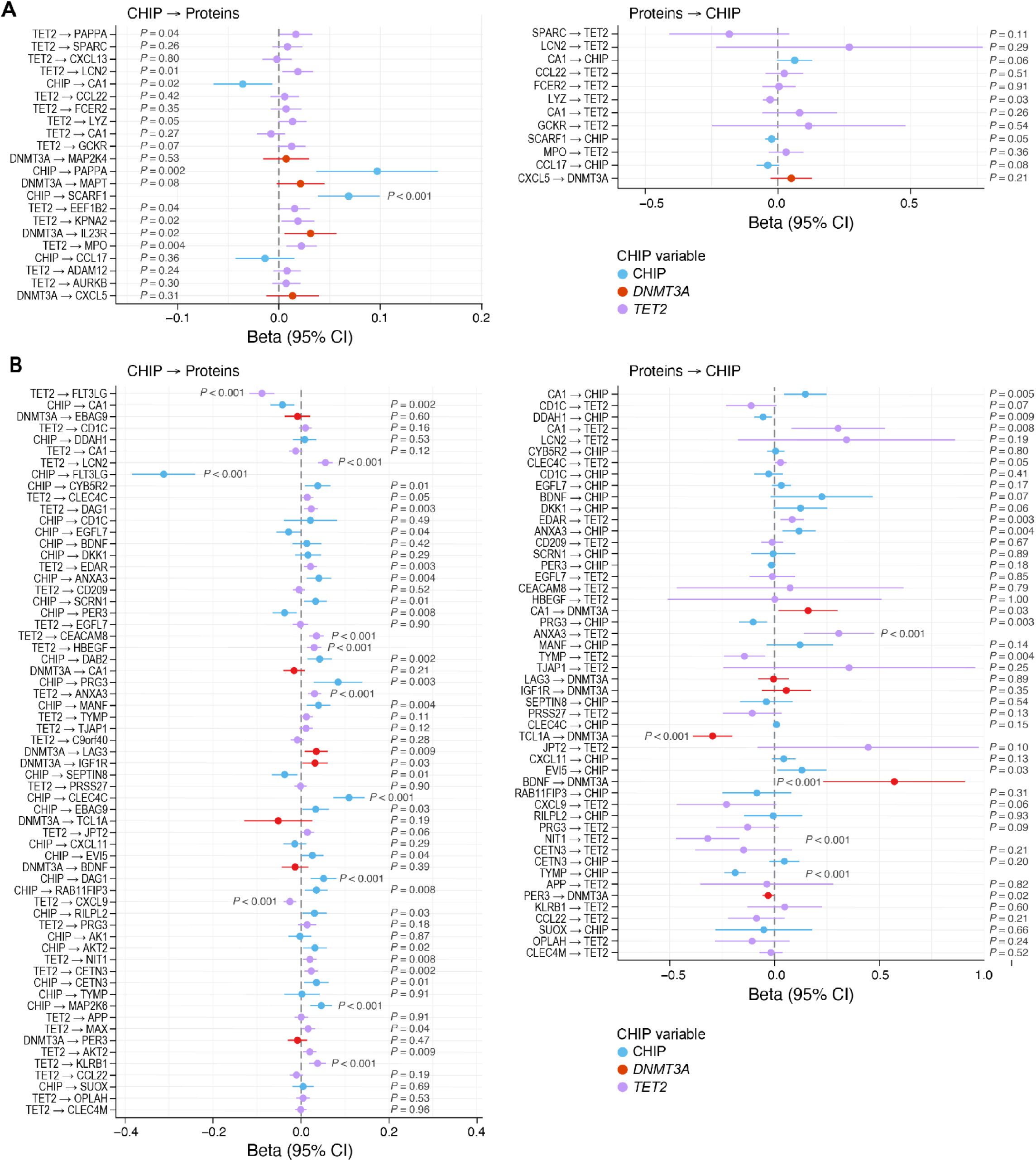
Estimation of bi-directional genetic causal effects between CHIP mutations and associated proteins. A. Proteins measured by SomaScan in TOPMed cohorts. B. Proteins measured by Olink in UK Biobank. For both A and B, we examined CHIP mutations’ genetic causal effects on proteins and proteins’ at FDR=0.05 level were examined. CHIP mutations were limited to overall CHIP, *DNMT3A*, and *TET2* given the availability of GWAS. Some proteins were not examined as no valid instruments were available. Inverse-variance weighted Mendelian randomization approach were used for the analysis.

Among the nine significant pairs from the TOPMed Somascan analysis, proteins from four pairs were also present in the UK Biobank Olink data. Two pairs (composite CHIP to CA1 and *TET2* to LCN2) showed consistent significance and causal directions in both datasets, while the other two pairs were not significantly associated and thus not included in MR analysis. It is important to note that association effects can encompass bidirectional causal influences. For instance, while *TET2* negatively associates with LCN2, the average causal direction shows that *TET2* positively influences LCN2 levels.

### Murine Evidence Corroborating Human Causal Discoveries

After showing consistency across two proteomics platforms with support for causality from Mendelian randomization, we examined the plasma levels of proteins influenced by *TET2* in 8–9-week-old mice with *Tet2* deletion in hematopoietic cells. Specifically, we selected LCN2 (significantly causal by *TET2* in both TOPMed cohorts and UK Biobank), MPO (significantly causal by *TET2* in TOPMed cohorts), as well as FLT3LG (significantly causal by *TET2* in UK Biobank) for enzyme-linked immunosorbent assay (ELISA) analysis in male and female with hematopoietic *Tet2* deficiency and WT mice given their disease relevance based on epidemiological evidence. Consistent with human genetic causal evidence, we found that hematopoietic *Tet2-/-* significantly increased plasma MPO levels in both male and female mice compared to WT mice, and male mice with hematopoietic *Tet2*-/- exhibited higher plasma levels of LCN2 compared to WT controls. However, though slightly decreased in hematopoietic *Tet2*-/- female mice, consistent with the causal direction in human genetics analysis, FLT3LG is not significantly different between hematopoietic *Tet2*-/- mice and control mice in both males and females (**Figure 6**).

**Figure 6.**
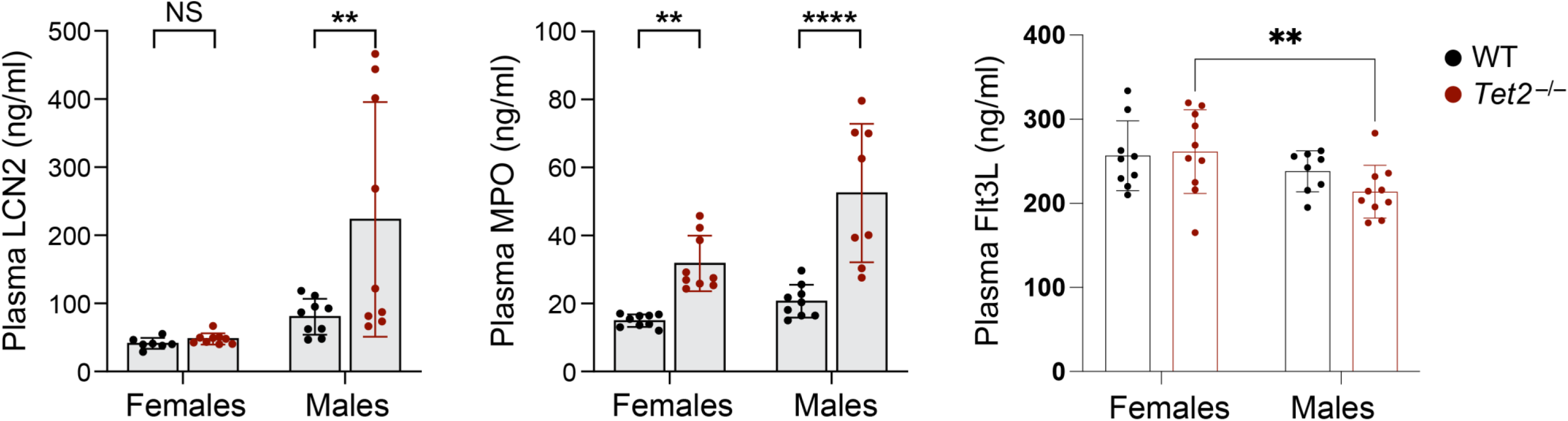
ELISA results of *Tet2^-/-^* and WT mice for selected plasma proteins whose level changes are associated with and causal by *TET2* in human. A. A protein of which the causal role of *TET2* is supported in both SomaScan and Olink. B. A protein of which the causal role of *TET2* is supported in SomaScan only. C. Proteins of which the causal role of *TET2* is supported in Olink only. Flt3L: FMS-related tyrosine kinase 3 ligand; LCN: Lipocalin 2; MPO: Myeloperoxidase; WT: wild-type

### Enriched Biological Pathways and Protein Networks

As SomaScan-measured proteins have wide analytic breadth across the proteome and are implicated in diverse pathways, we performed pathway analyses to investigate biological processes and regulatory mechanisms linked to the collective function of proteins associated with each CHIP driver gene from TOPMed cohorts’ results. The examined CHIP driver genes were broadly enriched in immune response and inflammation-related pathways and disease processes (**Supplemental Figures 14-17**). However, the significantly modulated pathways of different driver genes were involved in different immune activities and exhibited divergent effects. In addition to activating the cardiac hypertrophy signaling pathway that aligns with the observations of *DNMT3A*-mediated CH in heart failure^17^, *DNMT3A*-associated proteins activated pathways involved in acute responses to wound healing signaling and pathogen-induced cytokine storm signaling pathways^51^. In contrast, *TET2*-associated proteins modulated pathways implicated in autoimmunity and promoted chronic inflammation, activating the IL-17, STAT3, and IL-22 signaling pathways and inhibiting LXR/RXR activation. While *DNMT3A* and *TET2* appeared pro-inflammatory, *ASXL1* was linked to a number of reduced pro-inflammatory pathways, such as the STAT3 pathway that was predicted to be activated in the *TET2* pathway analysis and established IL-6 signaling pathway (**Figure 7**). We conducted sensitivity analysis where we limited to total quantified proteins as background for pathway analysis and yielded consistent results **(Supplemental Figures 18).** In secondary analyses, *JAK2-*associated proteins modulated tissue remodeling pathways, such as cardiac hypertrophy and pulmonary fibrosis idiopathic signaling pathways (**Supplemental Figures 19**).

**Figure 7.**
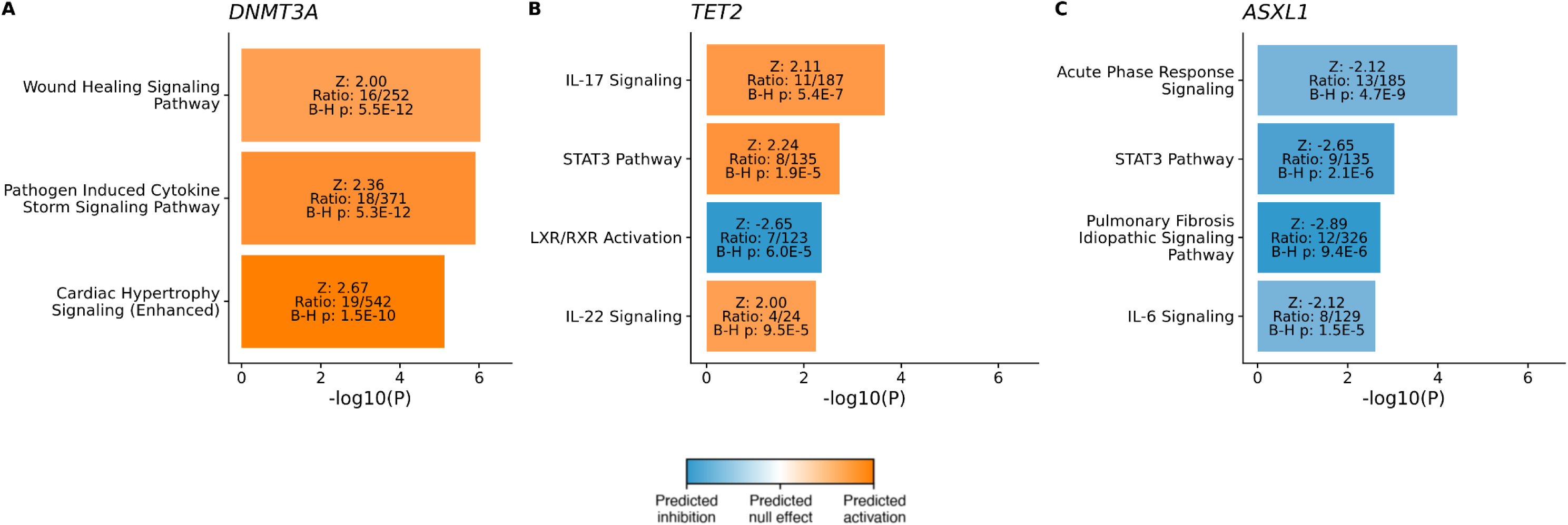
Significantly enriched and modulated pathways were identified among proteins associated with CHIP driver genes. Significantly enriched and modulated pathways corresponding to CHIP-associated proteins were derived based on known genetic and molecular relationships using IPA. The input was the Z-scores of the associations between major CHIP driver genes, i.e., *DNMT3A*, *TET2*, and *ASXL1*, and proteins that were significant at the P=0.05 level. The listed pathways fulfill two criteria: (1) within the top 30 most significantly enriched pathways by input proteins based on IPA analysis (P<0.05) and (2) being significantly modulated, either inhibited or activated, based on IPA analysis (Z>1.96). The orange indicates predicted activation, and the blue indicates predicted inhibition. The darker the color, the stronger the modulation effect. A. Significantly modulated canonical pathways implicated among proteins associated with *DNMT3A*. B. Significantly modulated canonical pathways implicated among proteins associated with *TET2*. C. Significantly modulated canonical pathways implicated among proteins associated with *ASXL1*. CHIP: Clonal hematopoiesis of indeterminate potential; IPA: Ingenuity Pathway Analysis

### Shared Proteomic Associations in CHIP and CAD

We next used SomaScan proteins to investigate the shared proteomic associations between CHIP mutations and CAD. Again, we used SomaScan results to facilitate potential novel discoveries. We analyzed the cross-sectional associations between prevalent CAD, which were assessed at the visits of blood draws to maintain temporal consistency with CHIP measurement, and proteomics. Top CAD-associated proteins include known CAD biomarkers, such as N-terminal pro-BNP (NT pro-BNP; FDR = 5.7×10^-13^), C-reactive protein (CRP; FDR = 2.7×10^-4^), and troponin I and T (FDR = 5.2×10^-3^ and 7.4×10^-3^, respectively), consistent with prior studies (**Supplemental Table 21**)^52, 53, 54, 55^. A total of 68 proteins were also associated with composite CHIP, *DNMT3A*, *ASXL1*, or *TET2* and CAD at the nominal significance threshold. These shared proteins have diverse functions, primarily enriched in inflammation and immune response pathways. Specifically, a number of the shared proteins were implicated in the regulation of IL- 1β/NLRP3/IL-6 pathways, including proteins belonging to TNFRSF (such as TNFRSF1B and TNFRSF10D), members or receptors of the IL-1 cytokine family (such as IL-36α, IL-1Ra, IL- 1RL1, and IL-1R2), and other chemokines (such as CXCL13 and CXCL10). Notably, *Cxcl13* and several genes encoding IL-1-related proteins have been shown to have increased expressed in *Tet2^−/−^* peritoneal macrophages exposed to lipopolysaccharide and interferon-γ in vitro^32^. Other important inflammatory proteins are implicated, such as protein S100-A9, myeloperoxidase^56, 57^, as well as proteins validated in *Tet2^-/-^* Ldlr*^-/-^* mice that consumed an atherogenic diet, such as CXCL13 and lysozyme C. Consistent with the functions of proteins associated with CHIP, proteins related to both *TET2* and CAD are involved in the ECM, cell adhesion, and signaling, such as metalloproteinase inhibitor 3 (TIMP-3)^58^. Those associated with both *ASXL1* and CAD participate in enzyme and metabolism processes, such as proprotein convertase subtilisin/kexin type 9 (PCSK9). Proteins associated with both *DNMT3A* and CAD have diverse functions, with several proteins involved in the signaling and adhesion of neural cells, such as contactin-1^59, 60, 61^. Furthermore, 24 proteins were found to be associated with *JAK2*, including key proteins involved in hematopoietic traits, such as erythropoietin and ferritin (**Figure 8** and **Supplemental Table 22**).

**Figure 8.**
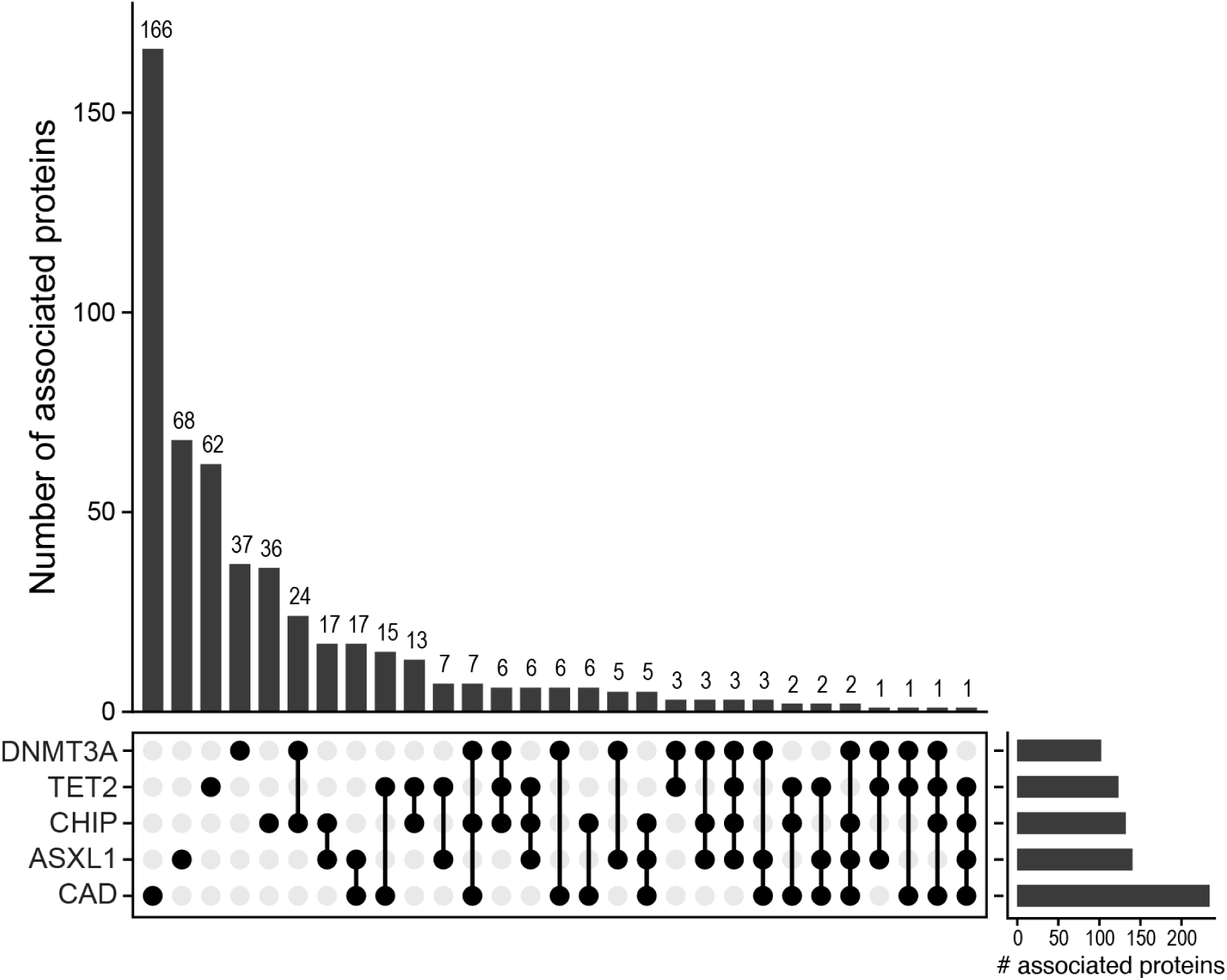
Upset plot showing overlapped and non-overlapped associated proteins between CHIP variables and CAD. A total of 68 proteins were associated with both prevalent CAD and any of the CHIP variables (composite CHIP, *DNMT3A, TET2*, and *ASXL1*) at P=0.05 level. For both CHIP variables and CAD, associations were assessed through linear regression models adjusting for age at sequencing, sex, race, batch (if applicable), type 2 diabetes status, smoker status, and the first ten principal components of genetic ancestry. PEER factors (the number of PEER factors varies by cohorts based on the sizes of study populations: 50 for JHS, MESA, and CHS; 70 for ARIC AA; 120 for ARIC EA) were adjusted in CHIP analysis only but not CAD analysis; this is because around 1/3 of them were associated with CAD, but in general not with CHIP. AA: African ancestry; ARIC: Atherosclerosis Risk in Communities; CAD: Coronary artery disease; CHIP, clonal hematopoiesis of indeterminate potential; CHS: Cardiovascular Heart Study; EA: European ancestry; FDR: False discovery rate; JHS: Jackson Heart Study; MESA: Multi-Ethnic Study of Atherosclerosis; PEER: Probabilistic estimation of expression residuals.

## Discussion

The various common mutations that drive CHIP yield differential associations with clinical outcomes^3, 10, 13^, and intermediate molecular phenotypes may facilitate a better understanding of these distinctions. Leveraging large-scale proteomics data from multiple diverse cohorts across two proteomics platforms, we identified and validated distinct proteomic signatures for common CHIP driver genes. Furthermore, our findings demonstrated the potential causal relationships between CHIP and those associated proteins through human genetics and murine ELISA experiments. More broadly, our study provides mechanistic insights offering novel CHIP driver gene-specific therapeutic strategies toward precision prevention of CHIP-associated clinical outcomes.

Despite convergence on clonal hematopoiesis^2, 3, 4^, mutations by CHIP driver genes had varied proteomic associations in both TOPMed cohorts and UK Biobank. Our study demonstrated that the three common driver genes linked to unique proteins with diverse functions, which subsequently aggregated into distinct pathways, suggesting these genes may associate with diseases through separate cellular pathways. For instance, although all examined driver mutations related to some proteins involved in immune responses, with particular enrichment in regulating IL-1β/NLRP3/IL-6 pathways, in line with existing knowledge that CHIP arising from several mutations induces an altered inflammatory state, different genes act differently. *DNMT3A*, the most prevalent CHIP driver gene, exhibited a relatively indolent profile with few and weak associations, primarily implicating first-line defense pathways. In contrast, *TET2*, associated with proteins predominantly involved in innate immunity, inflammation, and extracellular matrix remodeling, and *ASXL1*, enriched in proteins related to cell signaling and function and metabolic regulation, appeared to modulate chronic inflammation pathways, such as STAT3 and IL-6 signaling pathways. *JAK2*, the least prevalent CHIP mutated gene examined, was associated with the largest number of proteins exhibiting the strongest associations, enriched in hematopoietic traits. In particular, our study contributed evidence in humans of the anti-inflammatory role of *ASXL1* CHIP mutations, with pathway analysis showing predicted reduction across a large number of pro-inflammatory pathways, in direct contrast with other examined CHIP driver genes. This finding corresponds with the observed coexistence of pro- and anti-inflammatory characteristics in both zebrafish and murine macrophages with *Asxl1* mutations, and similarly in humans with *ASXL1* CHIP mutations^62, 63^. Moreover, leveraging multi-ancestry data from both men and women, our study highlighted sex and race differences in CHIP-proteomic associations, with the sex differences in CHIP’s impact on proteome being observed in our mice experiment as well. The factors driving differential associations across populations requires further investigation. Several of these findings agree closely with existing observations at the phenotypic level in humans or mice, substantiating the validity of our results and offering potential molecular mechanisms to explain these differential phenotypic associations^13, 64, 65^.

Association effects can encompass bidirectional causal influences when studying the relationship between CHIP and proteomics, both of which are dynamic. We conducted MR in our human study and ELISA in mice models to disentangle and validate the causal relationship between the two. We demonstrated that proteins with strong human causal evidence, such as *TET2* to MPO and LCN2, exhibit alignment in controlled murine models, supporting the validity of our human findings.

By examining shared proteomic associations between CHIP and CAD, in addition to proteins implicated in regulating the established CHIP-related IL-1β/NLRP3/IL-6 pathways, we implicate new potential associations. For instance, both *TET2* and CAD were linked to ECM-related proteins, such as TIMP3, an ECM-bound protein inhibiting a broad range of substrates, including matrix metalloproteinase^66, 67^. It is a potent angiogenesis inhibitor^68^ and has been shown to play crucial roles in cardiac remodeling and cardiomyopathy^69^. Recent studies also suggest its therapeutic potential for heart failure^70^. Similarly, *ASXL1* and CAD were associated with metabolism-related proteins like PCSK9, a key regulator of lipid metabolism. Inhibitors targeting PCSK9 lower LDL levels and reduce the risk of cardiovascular diseases^71^. Furthermore, *DNMT3A* and CAD shared associations with signaling and adhesion proteins in neural cells, such as contactin-1. This cell adhesion molecule plays a critical role in various aspects of neural development and function^59, 60, 61^, and has also been identified as a cardiac biomarker by several studies^72, 73^. Our study has limitations. Firstly, both molecular and environmental confounders might affect the associations between CHIP and the plasma proteome. Also, SomaScan proteomics from TOPMed cohorts was measured by two SomaScan platforms (1.3K and 5K), which may introduce heterogeneity due to technical differences. To address that, we adjusted for recognized confounders, applied PEER factors to mitigate technical noise, performed sensitivity analyses to reduce residual and unmeasured confounding as much as we could, and also reported heterogeneity p-value for meta-analysis. Secondly, our study primarily utilized linear models to explore associations between CHIP driver genes and proteins. However, there may be interactions and non-linear effects at play. To address this, we assessed interactions between CHIP variables and both sex and race for proteins, demonstrating differential associations in our stratified analysis. Thirdly, the cross-sectional design of our study constrains our ability to establish temporality and causality. Nevertheless, our experimental murine data provides some ability to infer associations resulting from *TET2* mutations.

These results, taken together, provide a comprehensive human plasma proteomic profile of clonal hematopoiesis. These novel findings inform us of the biological mechanism of various CHIP mutations and the potential development and testing of interventions to mitigate associated diseases observed in carriers of these mutations.

## Materials and Methods

### Human study participants

TOPMed is a research program generating genomic data from DNA isolated from blood and other omics data for more than 80 NHLBI-funded research studies with extensive phenotype data^22^. Our current study includes five community-based cohorts: JHS, MESA, CHS, ARIC, and UKB. The secondary use of data for this analysis was approved by the Massachusetts General Hospital Institutional Review Board (protocol 2016P001308 and protocol 2021P002228) and, for the UKB data, facilitated through UKB Application 7089.

JHS is a longitudinal cohort study of 5,306 self-identified Black men and women recruited in 2000-04 from Jackson, Mississippi^25^. Our study included 2,058 individuals with whole-genome sequencing (WGS) and plasma proteomic profiling data^22^. MESA is a multi-ancestry prospective cohort of 6,814 self-identified White, Black, Hispanic, or Asian (predominately of Chinese descent) men and women recruited in 2000-02^26^. We included 976 participants randomly selected for WGS and plasma proteomic profiling analysis^74^. CHS is a bi-ancestry (White and Black) longitudinal study of 5,888 men and women 65 years or older at recruitment (1989-90 or 1992-93)^27^. We analyzed data from 1,689 participants with WGS and plasma proteomics measurements who consented to genetics study^28^. ARIC is an ongoing longitudinal cohort of 15,792 middle-aged, mostly black and white participants recruited in 1987-89^75^. We included 8,188 participants with valid whole exome sequencing (WES) data and plasma proteomics measurements at Visit 2 or 3 in our study. These four cohorts, with data from a total of 12,911 participants, were used for the current analysis. UKB is a comprehensive, prospective cohort study consisting of approximately 500,000 men and women, predominantly of EA, who were aged between 40 and 69 years at the time of recruitment from 2006 to 2010 across the UK. Our analysis included a subset of 48,922 participants for whom both WES and plasma proteomic profiling data were available^23, 76^.

The mean age of the participants at the time of DNA sample collection was 57.4 years (55.8 years for JHS, 60.2 years for MESA, 73.3 years for CHS, 57.8 years for ARIC, and 56.8 years for UKB). Except for JHS, which includes only Black participants, all cohorts are bi-ancestry or multi-ancestry. All cohorts included participants of both sexes, with the percentage of men ranging from 39% to 47%.

### CHIP calling

WGS and CHIP calling in JHS, MESA, and CHS were previously performed and have been described in detail elsewhere^13^. The same procedure was applied for WES data in ARIC and UK Biobank^13, 19^. In brief, whole blood-derived DNA was sequenced at an average depth of 38× using Illumina HiSeq X Ten instruments. All sequences in CRAM files were remapped to the hs38DH 1000 Genomes build 38 human genome references, following the protocol published previously^77^. Single nucleotide polymorphisms (SNPs) and short indels were jointly discovered and genotyped across the TOPMed samples using the GotCloud pipeline ^78^. CHIP mutations were identified using Mutect2 software if one or more of a prespecified list of pathogenic somatic variants in 74 genes that drive clonal hematopoiesis and myeloid malignancies were present^13, 79^. A Panel of Normals (PON) minimized sequencing artifacts and Genome Aggregation Database (gnomAD) filtered likely germline variants from the putative somatic mutations call set^80^. Each Variant Call Format (VCF) file was annotated using ANNOVAR software^81^. Variants were retained for further curation if they met the following criteria: total depth of coverage ≥ 10, number of reads supporting the alternate allele ≥ 3, ≥ 1 read in both forward and reverse direction supporting the alternate allele, and variant allele fraction (VAF) ≥ 0.02. In particular, variants with VAF ≥ 0.1 were defined as expanded CHIP clones.

### Human proteomic measurements

The relative concentrations of plasma proteins or protein complexes from the blood samples of JHS, MESA, CHS, and ARIC were measured by the SomaScan (SomaLogic; Boulder, CO) using an aptamer (SOMAmer)-based approach, while proteomics of WHI was measured by Olink (Olink Proteomics; Uppsala, Sweden) using a proximity-extension immunoassay-based method. Detailed information on these technologies can be found in the corresponding manufacturer’s protocols^24, 82^. JHS and MESA utilized a 1.3K human protein platform, while CHS and ARIC used a 5K human protein platform. We focused on 1,148 proteins shared by both SomaScan platforms. Protein measurements were reported as relative fluorescence units (RFUs)^21^. There were no missing values in the SomaScan proteomic data, and details of the quality control of the proteins were described elsewhere^83^. Proteomics measured by Olink in UK Biobank included 2,917 proteins from cardiovascular, inflammation, cardiometabolic, neurology, oncology, and other panels. Proteins with ≥ 10% missingness were excluded. Participants who had >10% missing proteomics data were excluded. When examining the associations between CHIP variables and proteomics, we did not further process the missingness. When generating PEER factors for proteomics data, which requires complete data, we used K-nearest neighbors imputation^84, 85^. In this study, all proteins underwent log2 transformation. UK Biobank proteins were additionally normalized centrally.

### Regression model and meta-analysis for human analysis

We examined the cross-sectional associations between CHIP mutations and 1,148 plasma proteins measured by SomaScan within four TOPMed cohorts, which were then meta-analyzed, and between CHIP mutations and 2,917 plasma proteins measured by Olink in UK Biobank. TOPMed cohorts and UK Biobank analysis were conducted separately, given the limited overlapping in proteins between the two platforms. CHIP was modeled both as a composite and individually for the most common or pathogenic drivers (*DNMT3A*, *TET2*, *ASXL1,* and *JAK2*), using conventional thresholds for all mutations (VAF ≥2%) and expanded forms (VAF ≥10%), resulting in a total of ten CHIP mutations. Consistent with a recent analysis^31^, ARIC was separated into two subcohorts: ARIC EA and ARIC AA. To reduce redundant information and multiple testing burdens, we collapsed the composite CHIP and each driver gene with their corresponding expanded forms, retaining the one with stronger associations (larger absolute Z score). Within each cohort or subcohort (ARIC), linear regression models were fitted with CHIP mutations as exposures, log-transformed proteins as outcomes, and various covariates, including age at sequencing, sex, race, batch, center, diagnoses of type 2 diabetes mellitus at the time of enrollment, ever-smoker status, first ten PCs of genetic ancestry, and PEER factors as covariates. PEER factors were adjusted to account for hidden confounders, such as batches, that may influence clusters of proteins^86^. The number of PEER factors varied by cohorts based on study population size: 50 for JHS, MESA, and CHS; 70 for ARIC AA; 120 for ARIC EA^31, 87^; and 150 for UK Biobank. *JAK2* analyses were only conducted in cohorts with more than 5 participants with JAK2 mutations (i.e., CHS, ARIC EA, and UK Biobank only) to optimize power and generate more reliable effect estimates; given the required subsetting; *JAK2* analyses were considered secondary analyses. Next, linear regression results from each discovery cohort were meta-analyzed using inverse-variance weighted fixed-effect meta-analysis. We conducted stratified analysis by sex (female and male) and by race (Black and White; TOPMed cohorts only). For CHIP variable-protein pairs that are only significant in either male or female stratified analysis, we introduced and tested interaction terms between the corresponding CHIP variables and sex across all discovery cohorts. Likewise, for pairs that are only significant in Black or White stratified analysis, we tested for interaction terms between the CHIP variables and race in the combined analysis of ARIC, as ARIC has a good number of and relatively balanced White and Black participants. We also conducted secondary analyses additionally adjusting for eGFR or blood cells (platelet and WBC) in ARIC^88^. Linear regression models were performed using R function ‘glm’ while fixed-effects meta-analysis and tests of heterogeneity were conducted using the R package ‘meta’. We controlled the FDR using the Benjamini-Hochberg procedure and set an FDR threshold of 0.05 for significance.

### Cross-platform replication for human proteomic associations

We assessed the compatibility of associations between CHIP mutations and proteomic data measured using two highly multiplexed technologies for large-scale proteomics measurements: aptamer-based (SomaScan 1.3K) and proximity-extension immunoassay (Olink 3K) platforms^21, 24^. A total of 493 were overlapped between the two platforms as matched by UniProt IDs. We compared our results of these proteins between meta-analysis results from TOPMed cohorts based on SomaScan-measured proteins and results from UK Biobank based on Olink-measured proteins, with assessing shared CHIP variable-proteomic pairs between both groups.

### Mendelian randomization analyses

We performed two-sample MR analyses for CHIP-proteomic pairs with FDR < 0.05 to estimate the causal effects of CHIP on proteomics and vice versa. CHIP variables being tested include composite CHIP, *DNMT3A*, and *TET2*, given GWAS availability. The GWAS summary statistics for CHIP were from our CHIP GWAS meta-analysis for 648,992 multi-ancestry individuals in the UK Biobank, TOPMed, Vanderbilt BioVU, and Mass General Brigham Biobank^89^. For genetic instruments of proteins, we obtained SomaScan pQTL data from 35,892 Icelanders^49^ and Olink pQTL data from 48,922 UK Biobank-Pharma Proteomics Project participants who had their circulating proteomes profiled. All GWAS summary statistics assumed an additive genetic model. We used the inverse-variance-weighted (IVW) method for genetic instruments with more than one cis-pQTL and the Wald ratio estimator for instruments with only one cis-pQTL. IVW estimates were adjusted for residual correlation between genetic variants.

### Mouse experiments

All experiments were approved by the Institutional Animal Care and Use Committees and were conducted in accordance with the guidelines of the American Association for Accreditation of Laboratory Animal Care and the National Institutes of Health. To create animals with specific *Tet2* deletion in hematopoietic cells, we crossed *Tet2*-floxed line B6;129S-*Tet2*^tm1.1Iaai^/J (Jax Cat. No. 017573) with mice bearing constitutive expression of Cre recombinase under control of the *Vav1* promoter (B6.Cg-Commd10^Tg(Vav1-icre)A2Kio^/J; Jax Cat. No. 008610).

At 8-9 weeks old, vav1-cre; *Tet2-/-* (*Tet2-/-*) and vav1-cre; WT (WT) mice were sacrificed by CO2 euthanasia and blood was collected through cardiac puncture.

### ELISA

Mouse plasma levels of MPO, LCN2, and FLT3LG were quantified by ELISA following the manufacturer’s guidelines (Abcam, cat#275109; R&D systems cat# DY1857 and #DY427).

### Pathway analysis for human proteomic association results

We conducted pathway analyses of proteins associated with each CHIP driver gene at a P=0.05 threshold. We applied a nominal threshold (P=0.05) for selecting proteomic associations for pathway analysis, ensuring a comprehensive view of pathway enrichment by including a wide range of associated proteins. We input the sets of Z-scores of 102 *DNMT3A*-associated proteins, 123 *TET2*-associated proteins, and 140 *ASXL1*-associated proteins, which were organized into canonical pathways by the Ingenuity Pathway Analysis (IPA) tool. IPA pathways were constructed within the Ingenuity Knowledge Base, a large structured collection of findings containing nearly 5 million entries manually curated from the biomedical literature or integrated from third-party databases^90^. The network comprises ∼40,000 nodes connected by ∼1,480,000 edges representing experimentally observed cause-effect relationships related to expression, transcription, activation, molecular modification, transportation, and binding events. IPA utilizes a right-tailed Fisher’s exact test to evaluate the enrichment of CHIP driver genes-associated proteins in each pathway, as well as to infer their potential cause-effect relationships.

### Human coronary artery disease ascertainment

The four TOPMed cohorts have conducted active surveillance for coronary artery disease (CAD) events through annual follow-up by phone calls, surveys, and/or interviews, and abstracting medical records, hospitalization records, and death certificates^91, 92, 93, 94, 95^. In JHS, CAD was defined as myocardial infarction (MI), death due to CAD, or cardiac procedures, including percutaneous transluminal coronary angioplasty, stent placement, coronary artery bypass grafting, or other coronary revascularization^91^. In MESA, CAD included MI, death due to CAD, resuscitated cardiac arrest, and revascularization^92^. In CHS, CAD included MI, death due to CAD, angina pectoris, and cardiac procedures, including angioplasty and coronary artery bypass graft^96^. In ARIC, CAD included MI observed on ECG, self-reported doctor-diagnosed heart attack, or self-reported cardiovascular surgery or coronary angioplasty, as well as study-adjudicated CAD cases between visit 1 and visit 2 or 3^97^. In our study, we defined prevalent CAD cases as those occurring before the blood sample collection visit, where CHIP measurements were also taken. By aligning the time points for prevalent CAD and CHIP assessments, we ensured a fair comparison between them in later analyses.

### Shared proteomic associations between CHIP and CAD in human

We investigated the shared proteomic associations between CHIP mutations and CAD using four discovery TOPMed cohorts (N=12,911), JHS, MESA, CHS, and ARIC, same study population for examining the associations between CHIP mutations and proteomics in the main analysis. We first examined the cross-sectional associations between prevalent CAD, assessed at the same visits as blood draws to maintain temporal consistency with CHIP measurements, and proteomics. For the associations between CAD and proteomics, we employed the same linear models used to study the associations between CHIP mutations and proteomics. Specifically, we again used processed proteomics as the outcome, replaced CHIP with CAD as the exposure, and adjusted for the same set of covariates excluding PEER factors (age at sequencing, sex, race, batch, center, type 2 diabetes mellitus diagnoses at enrollment, ever-smoker status, and the first ten PCs of genetic ancestry). Our decision not to adjust for PEER factors in the CAD and proteomics association analysis was based on a previous report indicating that PEER factors can capture proteomic variation related to disease mechanisms^98^. Additionally, empirical evidence from our own analysis showed that several PEER factors were associated with CAD, which could remove relevant signals. However, PEER factors were generally not associated with CHIP mutations, and with or without adjusting for PEER factors yielded consistent associations between CHIP mutations and proteomics. Subsequently, we identified the intersection of proteins associated with both CHIP mutations and CAD ascertained at the same visits at a P=0.05 level. We then categorized these proteins according to the type of CHIP mutations and investigated any differential enrichment by distinct types of CHIP mutations.

## Data availability

TOPMed individual-level DNA and proteomics data used in this analysis are available through restricted access via the dbGaP. UK Biobank individual-level data are available for request by application (https://www.ukbiobank.ac.uk). All code used for the described analysis will be uploaded to GitHub once the manuscript is accepted for publication.

## Acknowledgments

Whole genome sequencing (WGS) for the Trans-Omics in Precision Medicine (TOPMed) program was supported by the National Heart, Lung and Blood Institute (NHLBI). WGS for “NHLBI TOPMed: Multi-Ethnic Study of Atherosclerosis (MESA)” (phs001416.v1.p1) was performed at the Broad Institute of MIT and Harvard (3U54HG003067-13S1). Centralized read mapping and genotype calling, along with variant quality metrics and filtering were provided by the TOPMed Informatics Research Center (3R01HL-117626-02S1). Phenotype harmonization, data management, sample-identity QC, and general study coordination, were provided by the TOPMed Data Coordinating Center (3R01HL-120393-02S1), and TOPMed MESA Multi-Omics (HHSN2682015000031/HSN26800004). The Atherosclerosis Risk in Communities study has been funded in whole or in part with Federal funds from the National Heart, Lung, and Blood Institute, National Institutes of Health, Department of Health and Human Services, under Contract nos. (75N92022D00001, 75N92022D00002, 75N92022D00003, 75N92022D00004, 75N92022D00005). The authors thank the staff and participants of the ARIC study for their important contributions. Funding support for “Building on GWAS for NHLBI-diseases: the U.S. CHARGE consortium” was provided by the NIH through the American Recovery and Reinvestment Act of 2009 (ARRA) (5RC2HL102419). Metabolomics measurements were sponsored by the National Human Genome Research Institute (3U01HG004402-02S1). SomaLogic Inc. conducted the SomaScan assays in exchange for use of ARIC data. This work was supported in part by NIH/NHLBI grant R01 HL134320. Cardiovascular Health Study: This research was supported by contracts HHSN268201200036C, HHSN268200800007C, HHSN268201800001C, N01HC55222, N01HC85079, N01HC85080, N01HC85081, N01HC85082, N01HC85083, N01HC85086, 75N92021D00006, and grants U01HL080295, HL105756, and U01HL130114 from the National Heart, Lung, and Blood Institute (NHLBI), with additional contribution from the National Institute of Neurological Disorders and Stroke (NINDS). Additional support was provided by R01AG023629 from the National Institute on Aging (NIA). A full list of principal CHS investigators and institutions can be found at CHS-NHLBI.org. The content is solely the responsibility of the authors and does not necessarily represent the official views of the National Institutes of Health. The Jackson Heart Study (JHS) is supported and conducted in collaboration with Jackson State University (HHSN268201800013I), Tougaloo College (HHSN268201800014I), the Mississippi State Department of Health (HHSN268201800015I) and the University of Mississippi Medical Center (HHSN268201800010I, HHSN268201800011I and HHSN268201800012I) contracts from the National Heart, Lung, and Blood Institute (NHLBI) and the National Institute on Minority Health and Health Disparities (NIMHD). The authors also wish to thank the staffs and participants of the JHS. The MESA projects are conducted and supported by the National Heart, Lung, and Blood Institute (NHLBI) in collaboration with MESA investigators. Support for the Multi-Ethnic Study of Atherosclerosis (MESA) projects are conducted and supported by the National Heart, Lung, and Blood Institute (NHLBI) in collaboration with MESA investigators. Support for MESA is provided by contracts 75N92020D00001, HHSN268201500003I, N01-HC-95159, 75N92020D00005, N01-HC-95160, 75N92020D00002, N01-HC-95161, 75N92020D00003, N01-HC-95162, 75N92020D00006, N01-HC-95163, 75N92020D00004, N01-HC-95164, 75N92020D00007, N01-HC-95165, N01-HC-95166, N01-HC-95167, N01-HC- 95168, N01-HC-95169, UL1-TR-000040, UL1-TR-001079, UL1-TR-001420, UL1TR001881, DK063491, and R01HL105756. The authors thank the other investigators, the staff, and the participants of the MESA study for their valuable contributions. A full list of participating MESA investigators and institutes can be found at http://www.mesa-nhlbi.org. The WHI program is funded by the National Heart, Lung, and Blood Institute, National Institutes of Health, U.S. Department of Health and Human Services through contracts HHSN268201600018C, HHSN268201600001C, HHSN268201600002C, HHSN268201600003C, and HHSN268201600004C. We used Perplexity AI (https://www.perplexity.ai/) as searching engine, ChatGPT4 (https://chat.openai.com/) for debugging code and for assisting with tedious tasks such as counting how many proteins are in each functional categories. We also thank Mrs. Leslie Gaffney from the Broad Research Communication Lab for her valuable assistance in improving the display items.

## References

1. Steensma DP, et al. Clonal hematopoiesis of indeterminate potential and its distinction from myelodysplastic syndromes. Blood 126, 9–16 (2015).

2. Xie M, et al. Age-related mutations associated with clonal hematopoietic expansion and malignancies. Nat Med 20, 1472–1478 (2014).

3. Genovese G, et al. Clonal hematopoiesis and blood-cancer risk inferred from blood DNA sequence. N Engl J Med 371, 2477–2487 (2014).

4. Jaiswal S, et al. Age-related clonal hematopoiesis associated with adverse outcomes. N Engl J Med 371, 2488–2498 (2014).

5. Kim PG, et al. Dnmt3a-mutated clonal hematopoiesis promotes osteoporosis. J Exp Med 218, (2021).

6. Yu B, et al. Supplemental Association of Clonal Hematopoiesis With Incident Heart Failure. J Am Coll Cardiol 78, 42–52 (2021).

7. Miller PG, et al. Association of clonal hematopoiesis with chronic obstructive pulmonary disease. Blood 139, 357–368 (2022).

8. Bhattacharya R, et al. Clonal Hematopoiesis Is Associated With Higher Risk of Stroke. Stroke 53, 788–797 (2022).

9. Jaiswal S, et al. Clonal Hematopoiesis and Risk of Atherosclerotic Cardiovascular Disease. N Engl J Med 377, 111–121 (2017).

10. Jaiswal S. Clonal hematopoiesis and nonhematologic disorders. Blood 136, 1606–1614 (2020).

11. Wong WJ, et al. Clonal haematopoiesis and risk of chronic liver disease. Nature, (2023).

12. Natarajan P, Jaiswal S, Kathiresan S. Clonal Hematopoiesis: Somatic Mutations in Blood Cells and Atherosclerosis. Circ Genom Precis Med 11, e001926 (2018).

13. Bick AG, et al. Inherited causes of clonal haematopoiesis in 97,691 whole genomes. Nature 586, 763–768 (2020).

14. Kar SP, et al. Genome-wide analyses of 200,453 individuals yield new insights into the causes and consequences of clonal hematopoiesis. Nat Genet 54, 1155–1166 (2022).

15. Uddin MM, et al. Germline genomic and phenomic landscape of clonal hematopoiesis in 323,112 individuals. medRxiv, 2022.2007.2029.22278015 (2022).

16. Abplanalp WT, Mas-Peiro S, Cremer S, John D, Dimmeler S, Zeiher AM. Association of Clonal Hematopoiesis of Indeterminate Potential With Inflammatory Gene Expression in Patients With Severe Degenerative Aortic Valve Stenosis or Chronic Postischemic Heart Failure. JAMA Cardiol 5, 1170–1175 (2020).

17. Abplanalp WT, et al. Clonal Hematopoiesis-Driver DNMT3A Mutations Alter Immune Cells in Heart Failure. Circ Res 128, 216–228 (2021).

18. Nachun D, et al. Clonal hematopoiesis associated with epigenetic aging and clinical outcomes. Aging Cell 20, e13366 (2021).

19. Uddin MDM, et al. Clonal hematopoiesis of indeterminate potential, DNA methylation, and risk for coronary artery disease. Nat Commun 13, 5350 (2022).

20. Pietzner M, et al. Mapping the proteo-genomic convergence of human diseases. Science 374, eabj1541 (2021).

21. Rohloff JC, et al. Nucleic Acid Ligands With Protein-like Side Chains: Modified Aptamers and Their Use as Diagnostic and Therapeutic Agents. Molecular therapy Nucleic acids 3, e201 (2014).

22. Taliun D, et al. Sequencing of 53,831 diverse genomes from the NHLBI TOPMed Program. Nature 590, 290–299 (2021).

23. Sun BB, et al. Genetic regulation of the human plasma proteome in 54,306 UK Biobank participants. bioRxiv, 2022.2006.2017.496443 (2022).

24. Assarsson E, et al. Homogenous 96-plex PEA immunoassay exhibiting high sensitivity, specificity, and excellent scalability. PLoS One 9, e95192 (2014).

25. Taylor HA, Jr., et al. Toward resolution of cardiovascular health disparities in African Americans: design and methods of the Jackson Heart Study. Ethn Dis 15, S6–4-17 (2005).

26. Bild DE, et al. Multi-Ethnic Study of Atherosclerosis: objectives and design. Am J Epidemiol 156, 871–881 (2002).

27. Fried LP, et al. The Cardiovascular Health Study: design and rationale. Ann Epidemiol 1, 263–276 (1991).

28. Austin TR, et al. Proteomics and Population Biology in the Cardiovascular Health Study (CHS): design of a study with mentored access and active data sharing. Eur J Epidemiol 37, 755–765 (2022).

29. Wright JD, et al. The ARIC (Atherosclerosis Risk In Communities) Study: JACC Focus Seminar 3/8. J Am Coll Cardiol 77, 2939–2959 (2021).

30. Kessler MD, et al. Common and rare variant associations with clonal haematopoiesis phenotypes. Nature 612, 301–309 (2022).

31. Zhang J, et al. Plasma proteome analyses in individuals of European and African ancestry identify cis-pQTLs and models for proteome-wide association studies. Nat Genet 54, 593–602 (2022).

32. Fuster JJ, et al. Clonal hematopoiesis associated with TET2 deficiency accelerates atherosclerosis development in mice. Science 355, 842–847 (2017).

33. Lin TM, Galbert SP, Kiefer D, Spellacy WN, Gall S. Characterization of four human pregnancy-associated plasma proteins. Am J Obstet Gynecol 118, 223–236 (1974).

34. Wald N, et al. First trimester concentrations of pregnancy associated plasma protein A and placental protein 14 in Down’s syndrome. Bmj 305, 28 (1992).

35. Kirkegaard I, Uldbjerg N, Oxvig C. Biology of pregnancy-associated plasma protein-A in relation to prenatal diagnostics: an overview. Acta Obstet Gynecol Scand 89, 1118–1125 (2010).

36. Smith GC, Stenhouse EJ, Crossley JA, Aitken DA, Cameron AD, Connor JM. Early-pregnancy origins of low birth weight. Nature 417, 916 (2002).

37. Brekken RA, Sage EH. SPARC, a matricellular protein: at the crossroads of cell-matrix communication. Matrix Biol 19, 816–827 (2001).

38. Bradshaw AD, Sage EH. SPARC, a matricellular protein that functions in cellular differentiation and tissue response to injury. J Clin Invest 107, 1049–1054 (2001).

39. Tai IT, Tang MJ. SPARC in cancer biology: its role in cancer progression and potential for therapy. Drug Resist Updat 11, 231–246 (2008).

40. Lindskog S. Structure and mechanism of carbonic anhydrase. Pharmacol Ther 74, 1–20 (1997).

41. Steppan CM, et al. The hormone resistin links obesity to diabetes. Nature 409, 307–312 (2001).

42. Jackson DG. Hyaluronan in the lymphatics: The key role of the hyaluronan receptor LYVE-1 in leucocyte trafficking. Matrix Biol 78-79, 219–235 (2019).

43. Lee JY, Spicer AP. Hyaluronan: a multifunctional, megaDalton, stealth molecule. Curr Opin Cell Biol 12, 581–586 (2000).

44. Quach ME, Li R. Structure-function of platelet glycoprotein Ib-IX. J Thromb Haemost 18, 3131–3141 (2020).

45. Furie B, Furie BC. Role of platelet P-selectin and microparticle PSGL-1 in thrombus formation. Trends Mol Med 10, 171–178 (2004).

46. Hobbs CM, et al. JAK2V617F leads to intrinsic changes in platelet formation and reactivity in a knock-in mouse model of essential thrombocythemia. Blood 122, 3787–3797 (2013).

47. Maslah N, et al. Revisiting Diagnostic performances of serum erythropoïetin level and JAK2 mutation for polycythemias: analysis of a cohort of 1090 patients with red cell mass measurement. Br J Haematol 196, 676–680 (2022).

48. Chin-Yee B, et al. Serum erythropoietin levels in 696 patients investigated for erythrocytosis with JAK2 mutation analysis. Am J Hematol 97, E150–e153 (2022).

49. Eldjarn GH, et al. Large-scale plasma proteomics comparisons through genetics and disease associations. Nature 622, 348–358 (2023).

50. Rooney MR, et al. Comparison of Proteomic Measurements Across Platforms in the Atherosclerosis Risk in Communities (ARIC) Study. Clin Chem 69, 68–79 (2023).

51. Fajgenbaum DC, June CH. Cytokine Storm. N Engl J Med 383, 2255–2273 (2020).

52. Thygesen K, et al. Fourth Universal Definition of Myocardial Infarction (2018). Circulation 138, e618–e651 (2018).

53. Roberts WL. CDC/AHA Workshop on Markers of Inflammation and Cardiovascular Disease: Application to Clinical and Public Health Practice: laboratory tests available to assess inflammation--performance and standardization: a background paper. Circulation 110, e572–576 (2004).

54. Pearson TA, et al. Markers of inflammation and cardiovascular disease: application to clinical and public health practice: A statement for healthcare professionals from the Centers for Disease Control and Prevention and the American Heart Association. Circulation 107, 499–511 (2003).

55. Atherton JJ, et al. National Heart Foundation of Australia and Cardiac Society of Australia and New Zealand: Guidelines for the Prevention, Detection, and Management of Heart Failure in Australia 2018. Heart Lung Circ 27, 1123–1208 (2018).

56. Arnhold J. The Dual Role of Myeloperoxidase in Immune Response. Int J Mol Sci 21, (2020).

57. Averill MM, Kerkhoff C, Bornfeldt KE. S100A8 and S100A9 in cardiovascular biology and disease. Arterioscler Thromb Vasc Biol 32, 223–229 (2012).

58. Cabral-Pacheco GA, et al. The Roles of Matrix Metalloproteinases and Their Inhibitors in Human Diseases. Int J Mol Sci 21, (2020).

59. Bizzoca A, et al. Transgenic mice expressing F3/contactin from the TAG-1 promoter exhibit developmentally regulated changes in the differentiation of cerebellar neurons. Development 130, 29–43 (2003).

60. Bizzoca A, et al. F3/Contactin acts as a modulator of neurogenesis during cerebral cortex development. Dev Biol 365, 133–151 (2012).

61. Hu QD, et al. F3/contactin acts as a functional ligand for Notch during oligodendrocyte maturation. Cell 115, 163–175 (2003).

62. Avagyan S, et al. Resistance to inflammation underlies enhanced fitness in clonal hematopoiesis. Science 374, 768–772 (2021).

63. Yu Z, et al. Genetic modification of inflammation and clonal hematopoiesis-associated cardiovascular risk. medRxiv, 2022.2012.2008.22283237 (2023).

64. Fujino T, Kitamura T. ASXL1 mutation in clonal hematopoiesis. Exp Hematol 83, 74–84 (2020).

65. Montagner S, et al. TET2 Regulates Mast Cell Differentiation and Proliferation through Catalytic and Non-catalytic Activities. Cell Rep 15, 1566–1579 (2016).

66. Galis ZS, Sukhova GK, Lark MW, Libby P. Increased expression of matrix metalloproteinases and matrix degrading activity in vulnerable regions of human atherosclerotic plaques. J Clin Invest 94, 2493–2503 (1994).

67. Fan D, Kassiri Z. Biology of Tissue Inhibitor of Metalloproteinase 3 (TIMP3), and Its Therapeutic Implications in Cardiovascular Pathology. Front Physiol 11, 661 (2020).

68. Qi JH, et al. A novel function for tissue inhibitor of metalloproteinases-3 (TIMP3): inhibition of angiogenesis by blockage of VEGF binding to VEGF receptor-2. Nat Med 9, 407–415 (2003).

69. Moore L, Fan D, Basu R, Kandalam V, Kassiri Z. Tissue inhibitor of metalloproteinases (TIMPs) in heart failure. Heart Fail Rev 17, 693–706 (2012).

70. Takawale A, et al. Myocardial overexpression of TIMP3 after myocardial infarction exerts beneficial effects by promoting angiogenesis and suppressing early proteolysis. Am J Physiol Heart Circ Physiol 313, H224–h236 (2017).

71. Sabatine MS, et al. Evolocumab and Clinical Outcomes in Patients with Cardiovascular Disease. N Engl J Med 376, 1713–1722 (2017).

72. Yin X, et al. Protein biomarkers of new-onset cardiovascular disease: prospective study from the systems approach to biomarker research in cardiovascular disease initiative. Arterioscler Thromb Vasc Biol 34, 939–945 (2014).

73. Lau ES, et al. Cardiovascular Biomarkers of Obesity and Overlap With Cardiometabolic Dysfunction. J Am Heart Assoc 10, e020215 (2021).

74. Liu Y, et al. Blood monocyte transcriptome and epigenome analyses reveal loci associated with human atherosclerosis. Nat Commun 8, 393 (2017).

75. ARIC Investigators. The Atherosclerosis Risk in Communities (ARIC) Study: design and objectives. Am J Epidemiol 129, 687–702 (1989).

76. Backman JD, et al. Exome sequencing and analysis of 454,787 UK Biobank participants. Nature 599, 628–634 (2021).

77. Regier AA, et al. Functional equivalence of genome sequencing analysis pipelines enables harmonized variant calling across human genetics projects. Nat Commun 9, 4038 (2018).

78. Jun G, Wing MK, Abecasis GR, Kang HM. An efficient and scalable analysis framework for variant extraction and refinement from population-scale DNA sequence data. Genome Res 25, 918–925 (2015).

79. Benjamin D, Sato T, Cibulskis K, Getz G, Stewart C, Lichtenstein L. Calling Somatic SNVs and Indels with Mutect2. bioRxiv, 861054 (2019).

80. Karczewski KJ, et al. The mutational constraint spectrum quantified from variation in 141,456 humans. Nature 581, 434–443 (2020).

81. Wang K, Li M, Hakonarson H. ANNOVAR: functional annotation of genetic variants from high-throughput sequencing data. Nucleic Acids Res 38, e164 (2010).

82. Gold L, et al. Aptamer-based multiplexed proteomic technology for biomarker discovery. PLoS One 5, e15004 (2010).

83. Sun BB, et al. Genomic atlas of the human plasma proteome. Nature 558, 73–79 (2018).

84. Gadd DA, et al. Blood protein assessment of leading incident diseases and mortality in the UK Biobank. Nat Aging 4, 939–948 (2024).

85. Troyanskaya O, et al. Missing value estimation methods for DNA microarrays. Bioinformatics 17, 520–525 (2001).

86. Stegle O, Parts L, Piipari M, Winn J, Durbin R. Using probabilistic estimation of expression residuals (PEER) to obtain increased power and interpretability of gene expression analyses. Nat Protoc 7, 500–507 (2012).

87. Battle A, Brown CD, Engelhardt BE, Montgomery SB. Genetic effects on gene expression across human tissues. Nature 550, 204–213 (2017).

88. Inker LA, et al. New Creatinine- and Cystatin C-Based Equations to Estimate GFR without Race. N Engl J Med 385, 1737–1749 (2021).

89. Uddin MM, et al. Germline genomic and phenomic landscape of clonal hematopoiesis in 323,112 individuals. medRxiv, 2022.2007.2029.22278015 (2022).

90. Krämer A, Green J, Pollard J, Jr., Tugendreich S. Causal analysis approaches in Ingenuity Pathway Analysis. Bioinformatics 30, 523–530 (2014).

91. Keku E, et al. Cardiovascular disease event classification in the Jackson Heart Study: methods and procedures. Ethn Dis 15, S6–62-70 (2005).

92. McClelland RL, et al. 10-Year Coronary Heart Disease Risk Prediction Using Coronary Artery Calcium and Traditional Risk Factors: Derivation in the MESA (Multi-Ethnic Study of Atherosclerosis) With Validation in the HNR (Heinz Nixdorf Recall) Study and the DHS (Dallas Heart Study). J Am Coll Cardiol 66, 1643–1653 (2015).

93. Newman AB, et al. Dementia and Alzheimer’s disease incidence in relationship to cardiovascular disease in the Cardiovascular Health Study cohort. J Am Geriatr Soc 53, 1101–1107 (2005).

94. Dawber TR, Meadors GF, Moore FE, Jr. Epidemiological approaches to heart disease: the Framingham Study. Am J Public Health Nations Health 41, 279–281 (1951).

95. Kannel WB, Feinleib M, McNamara PM, Garrison RJ, Castelli WP. An investigation of coronary heart disease in families. The Framingham offspring study. Am J Epidemiol 110, 281–290 (1979).

96. Psaty BM, et al. Study of Cardiovascular Health Outcomes in the Era of Claims Data: The Cardiovascular Health Study. Circulation 133, 156–164 (2016).

97. White AD, et al. Community surveillance of coronary heart disease in the Atherosclerosis Risk in Communities (ARIC) Study: methods and initial two years’ experience. J Clin Epidemiol 49, 223–233 (1996).

98. Walker KA, et al. Large-scale plasma proteomic analysis identifies proteins and pathways associated with dementia risk. Nature Aging 1, 473–489 (2021).

