## Supplemental Figures for "Human Plasma Proteomic Profile of Clonal Hematopoiesis"

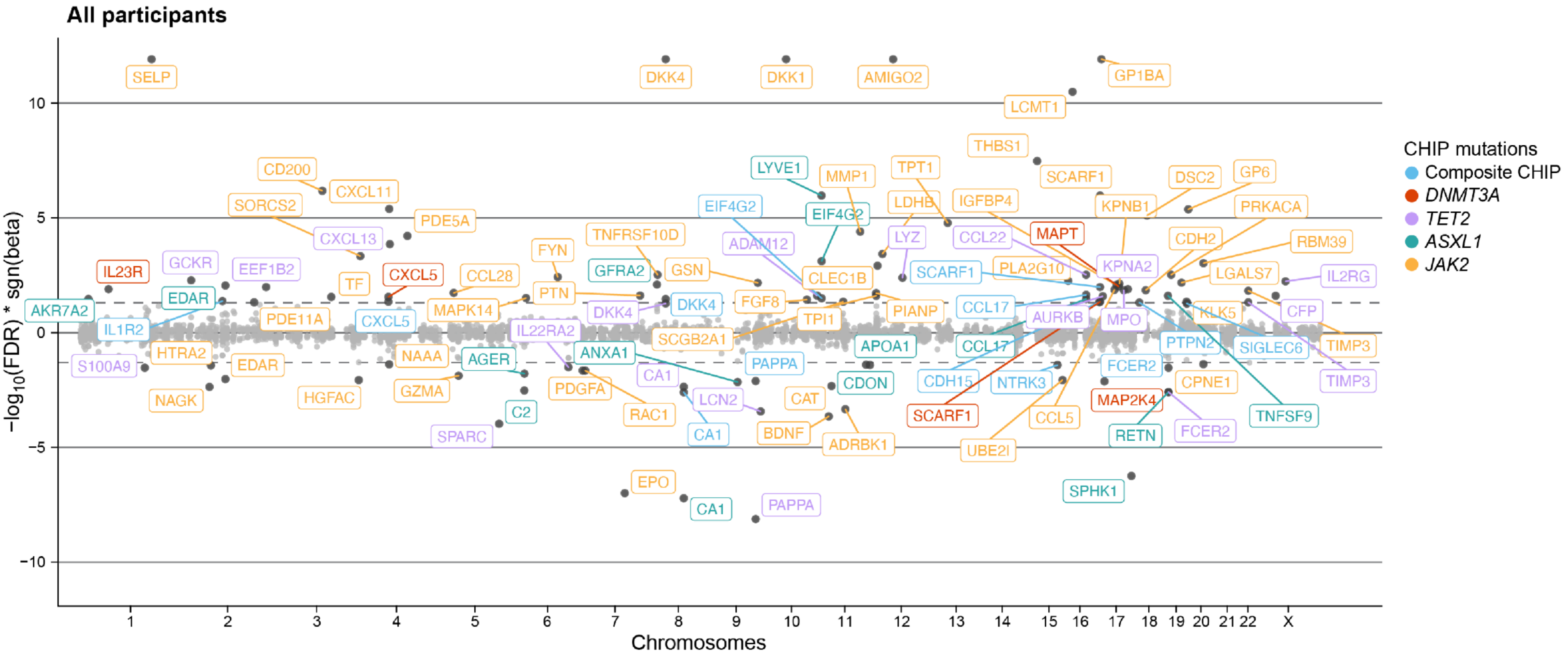


**Supplemental Figure 1. Meta-analyzed associations between CHIP mutations and circulating proteome among all participants in TOPMed cohorts** (N=12,911). Proteins that are associated at FDR=0.05 level (for 5,740 testings) are labeled with the corresponding SomaScan targets and colored in blue, red, green, orange, and purple, indicating significant associations with composite CHIP, DNMT3A, TET2, ASXL1, and JAK2, respectively. JAK2 analyses were only conducted in cohorts with greater than 5 participants with JAK2 mutations (i.e., CHS and ARIC EA). Associations were assessed through linear regression models adjusting for age at sequencing, sex, race, batch (if applicable), type 2 diabetes status, smoker status, first ten principal components of genetic ancestry, and PEER factors (the number of PEER factors varies by cohorts based on the sizes of study populations: 50 for JHS, MESA, and CHS; 70 for ARIC AA; 120 for ARIC EA). AA: African ancestry; ARIC: Atherosclerosis Risk in Communities; CHIP, clonal hematopoiesis of indeterminate potential; CHS: Cardiovascular Heart Study; EA: European ancestry; FDR: false discovery rate; JHS: Jackson Heart Study; MESA: Multi-Ethnic Study of Atherosclerosis; PEER: probabilistic estimation of expression residuals.

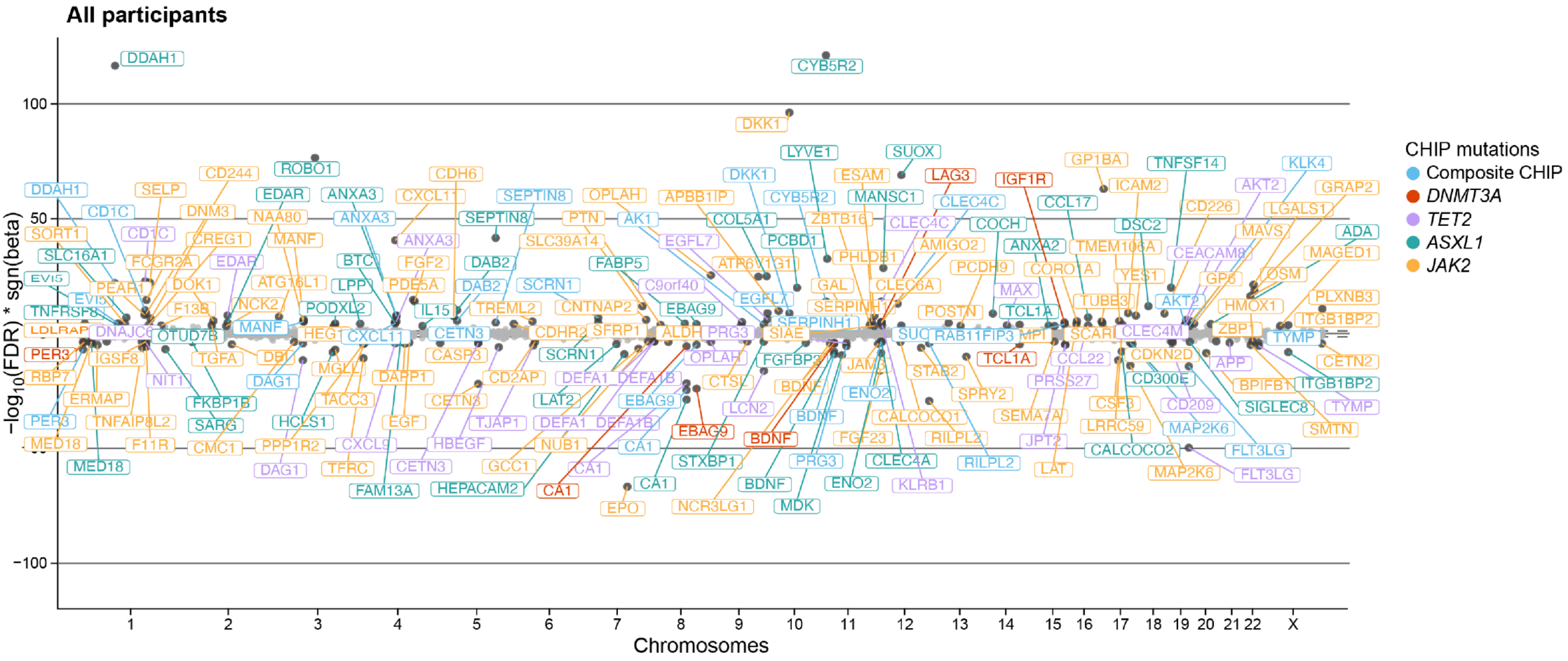


**Supplemental Figure 2. Associations between CHIP mutations and circulating proteome among all participants in UK Biobank (N=41,022).** Proteins that are associated at FDR=0.001 level (for 14,585 testings) are labeled with the corresponding Olink targets and colored in blue, red, purple, green, and orange, indicating significant associations with composite CHIP, *DNMT3A*, *TET2*, *ASXL1*, and *JAK2*, respectively. Associations were assessed through linear regression models adjusting for age at sequencing, sex, self-reported British White ancestry (if applicable), type 2 diabetes status, current smoker status, first ten principal components of genetic ancestry, and 150 PEER factors. CHIP: Clonal hematopoiesis of indeterminate potential; FDR: False discovery rate; PEER: Probabilistic estimation of expression residuals.

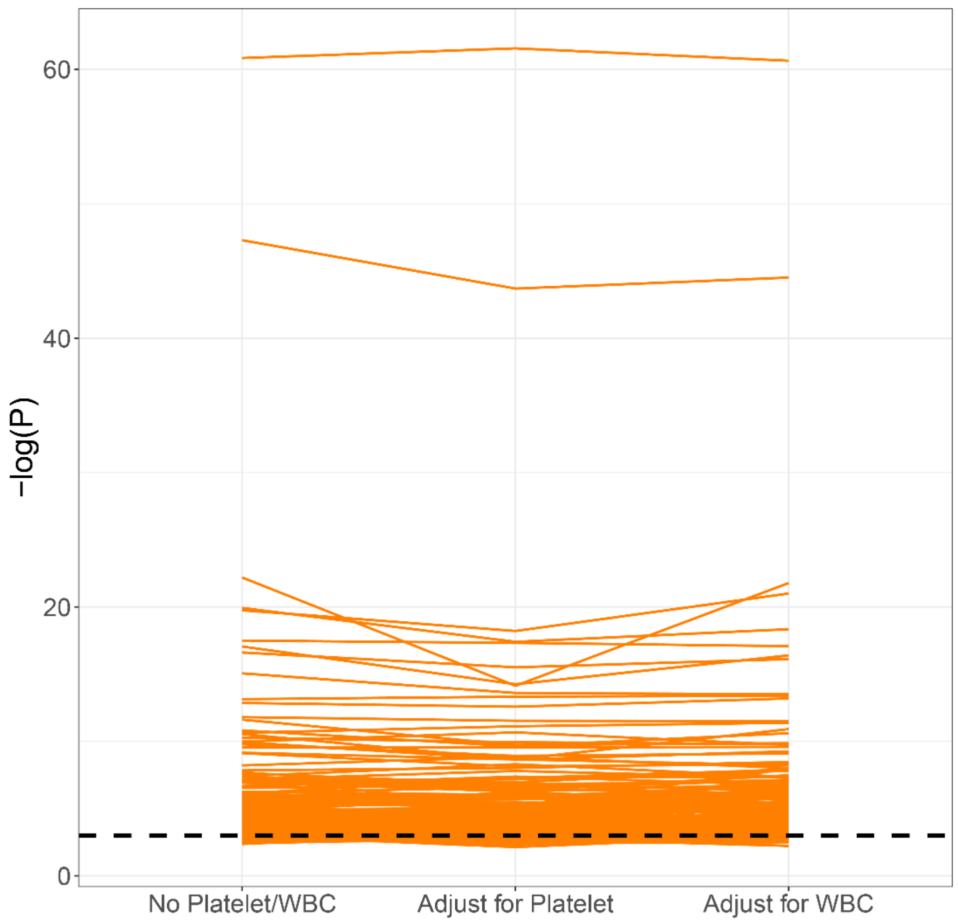


**Supplemental Figure 3. Proteomics associations with or without additionally adjusting for platelet or WBC in ARIC EA.** Here we present the *JAK2*-protein association changes with or without additionally adjusting for platelet or WBC. Associations were assessed through linear regression models adjusting for age at sequencing, sex, center, type 2 diabetes status, smoker status, first ten principal components of genetic ancestry, 120 PEER factors, and, for the second and third columns, platelet and WBC, respectively. The y-axis indicates the negative natural log of the P-value of those associations. ARIC: Atherosclerosis Risk in Communities; EA: European ancestry; WBC: white blood cell

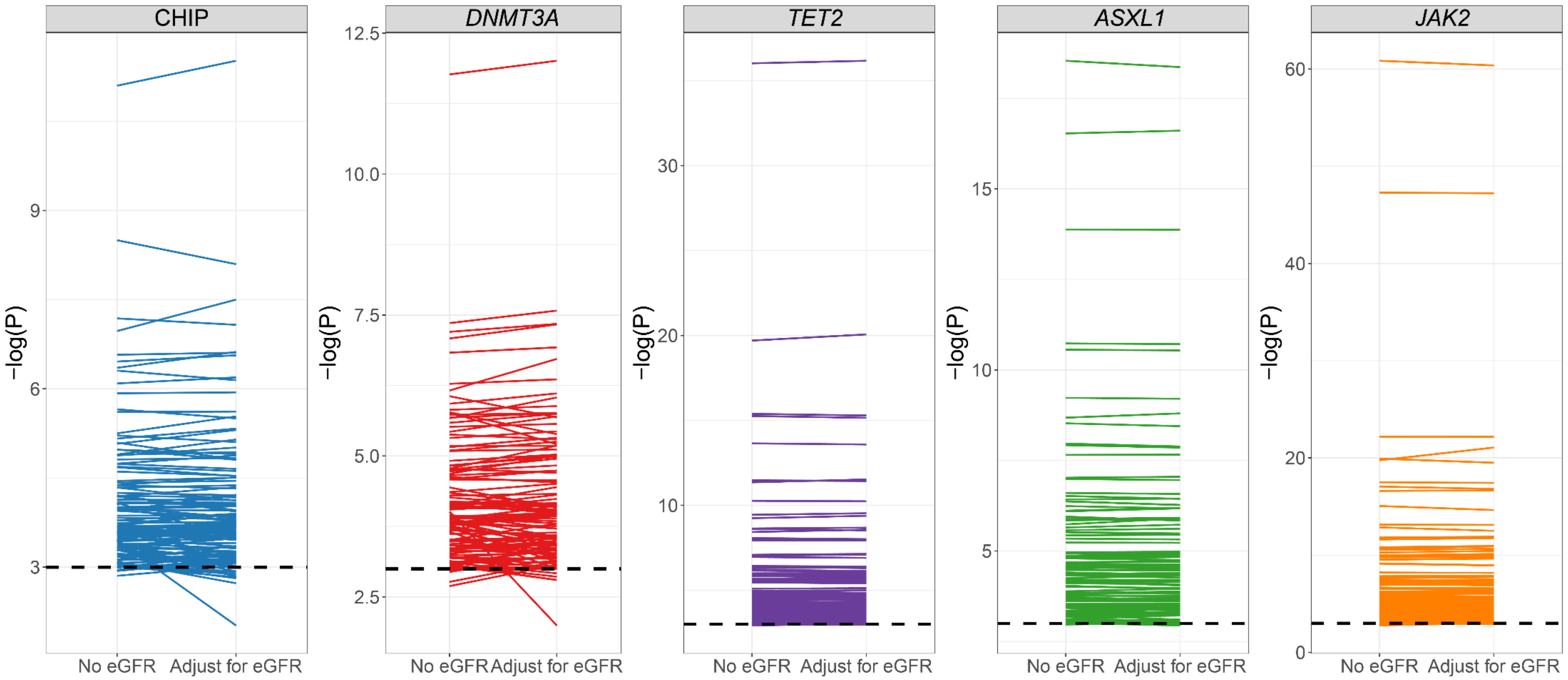


**Supplemental Figure 4. CHIP-proteomics associations with or without additionally adjusting for eGFR in ARIC EA.** Here we present the CHIP variable-protein association changes with or without additionally adjusting for eGFR for CHIP, *DNMT3A*, *TET2*, *ASXL1*, and *JAK2*. Associations were assessed through linear regression models adjusting for age at sequencing, sex, center, type 2 diabetes status, smoker status, first ten principal components of genetic ancestry, 120 PEER factors, and, for the second columns in each subplot, eGFR. The y-axis indicates the negative natural log of the P-value of those associations. ARIC: Atherosclerosis Risk in Communities; CHIP: Clonal hematopoiesis of indeterminate potential; EA: European ancestry; eGFR: estimated glomerular filtration rate.

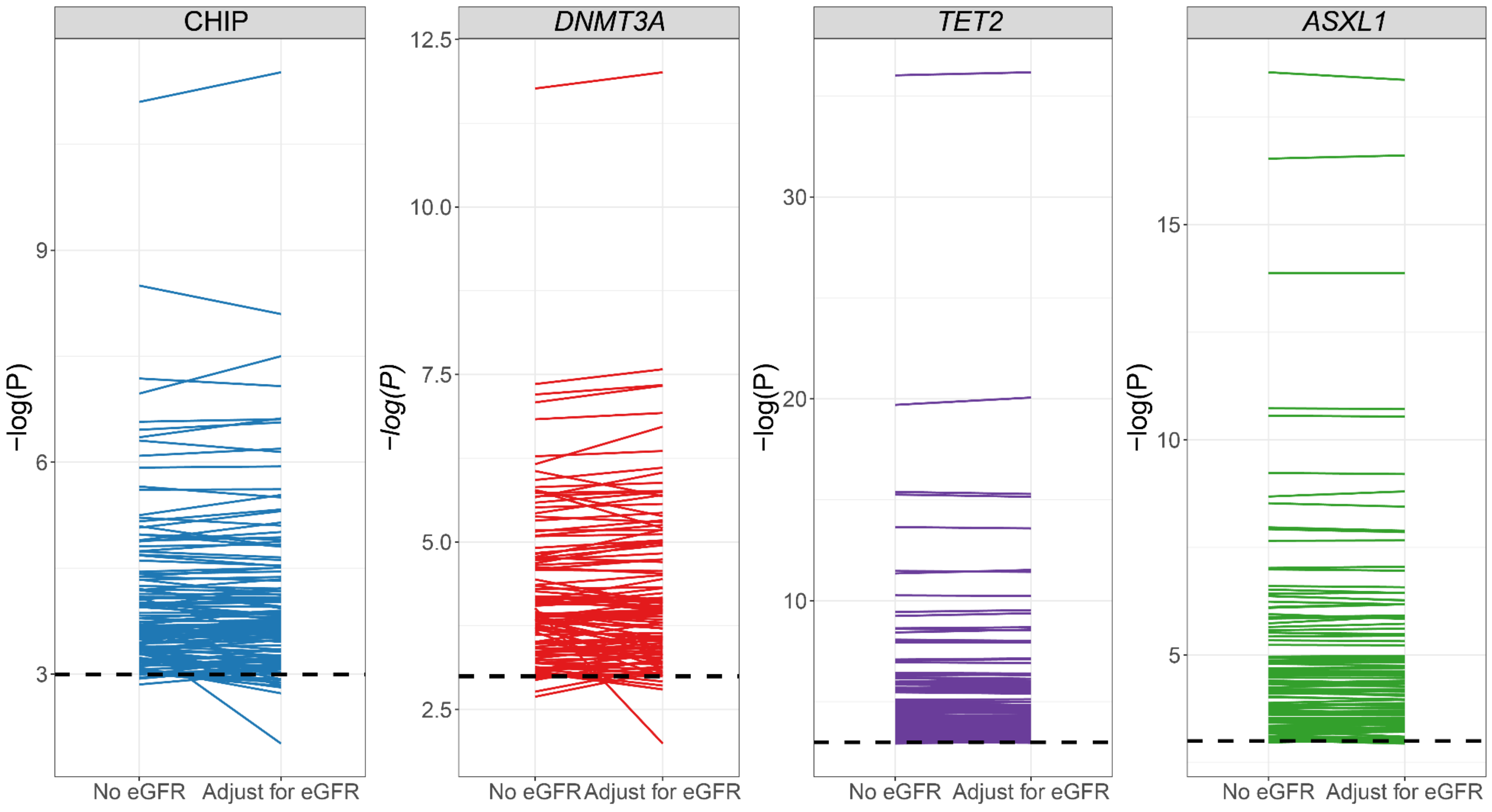


**Supplemental Figure 5. CHIP-proteomics associations with or without additionally adjusting for eGFR in ARIC AA.** Here we present the CHIP variable-protein associations changes with or without additionally adjusting for eGFR for CHIP, *DNMT3A*, *TET2*, and *ASXL1*. Associations were assessed through linear regression models adjusting for age at sequencing, sex, center, type 2 diabetes status, smoker status, first ten principal components of genetic ancestry, 120 PEER factors, and, for the second columns in each subplot, eGFR. The y-axis indicates the negative natural log of the P-value of those associations. AA: African Ancestry; ARIC: Atherosclerosis Risk in Communities; CHIP: Clonal hematopoiesis of indeterminate potential; eGFR: estimated glomerular filtration rate.

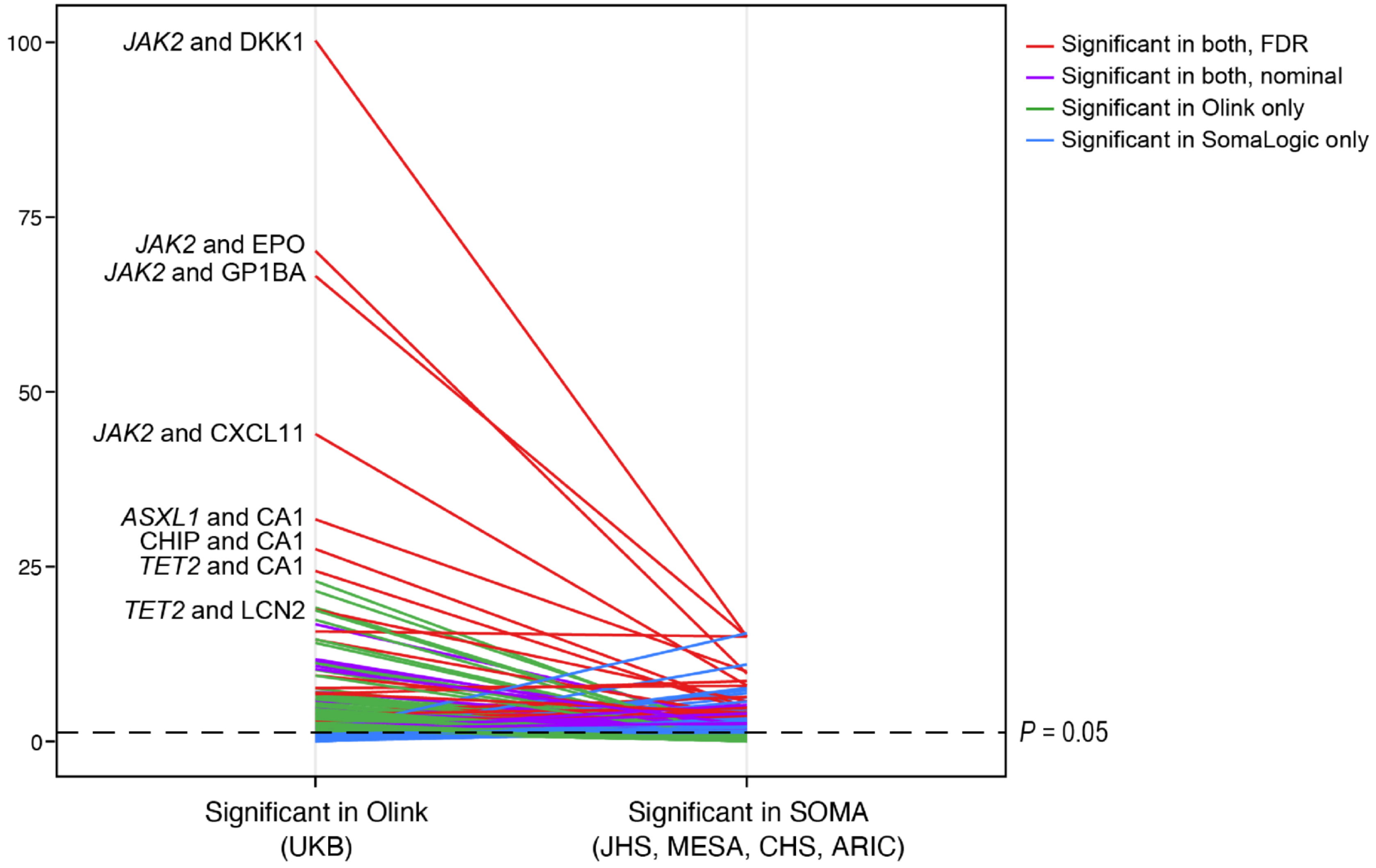


**Supplemental Figure 6. Cross-platform comparison of CHIP-proteomics associations.** A total of 493 proteins with unique UniPort ID were present in both Olink and SomaScan platforms, and here we present 373 of them that were significantly associated with at least one of the CHIP variables (composite CHIP, DNMT3A, TET2, and ASXL1) at P=0.05 in either UKB (Olink-based) or meta-analysis across female-only results from JHS, MESA, CHS, and ARIC (SomaScan-based). We examined the significantly associated CHIP variable-protein pairs' P-value differences between the Olink-based and SomaScan-based results. The y-axis indicates the negative log 10 of the P-value of those associations. The red color indicates CHIP variable-protein pairs significantly associated in both Olink-based and SomaScan-based results at FDR=0.05 level. The purple color indicates CHIP variable-protein pairs significantly associated in Olink-based and SomaScan-based results at P=0.05. The green color indicates CHIP variable-protein pairs significantly associated in only Olink-based but not SomaScan-based results. The blue color indicates CHIP variable-protein pairs significantly associated in only SomaScan-based but not Olink-based results. A total of 114 CHIP variable-protein pairs were significantly associated in both UKB and meta-analysis at P=0.05 threshold, and 26 of them were significant at FDR=0.05 level. ARIC: Atherosclerosis Risk in Communities; CHIP, clonal hematopoiesis of indeterminate potential; CHS: Cardiovascular Heart Study; JHS: Jackson Heart Study; MESA: Multi-Ethnic Study of Atherosclerosis; UKB: UK Biobank.

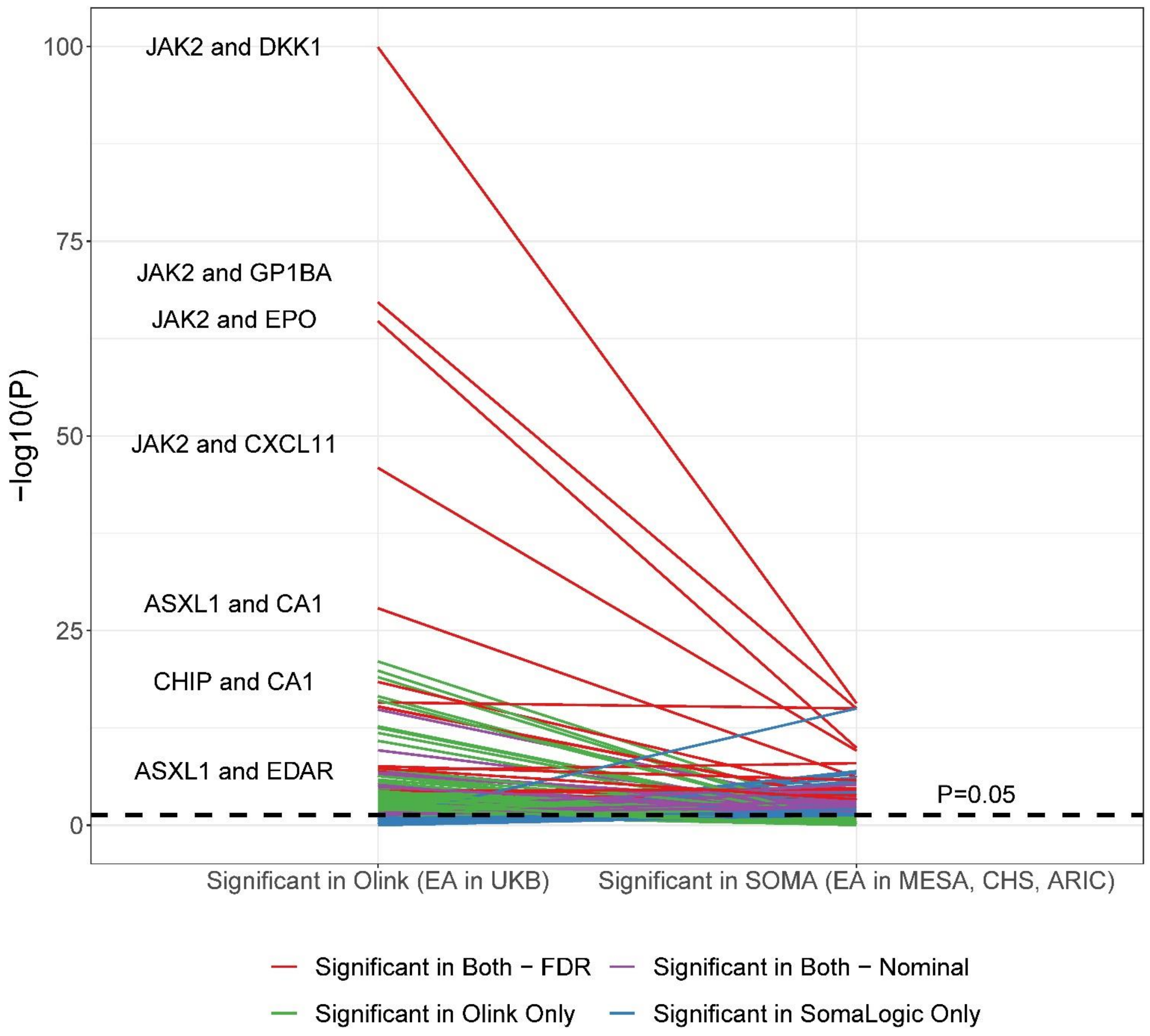


**Supplemental Figure 7. Cross-platform comparison of CHIP-proteomics associations among White participants only.** A total of 493 proteins with unique UniPort ID were present in both Olink and SomaScan platforms, and here we present 353 of them that were significantly associated with at least one of the CHIP variables (composite CHIP, DNMT3A, TET2, and ASXL1) at P=0.05 in either EA in UKB (Olink-based) or meta-analysis across female-only results from EA in MESA, CHS, and ARIC (SomaScan-based). We examined the significantly associated CHIP variable-protein pairs' P-value differences between the Olink-based and SomaScan-based results. The y-axis indicates the negative log 10 of the P-value of those associations. The red color indicates CHIP variable-protein pairs significantly associated in both Olink-based and SomaScan-based results at FDR=0.05 level. The purple color indicates CHIP variable-protein pairs significantly associated in Olink-based and SomaScan-based results at P=0.05. The green color indicates CHIP variable-protein pairs significantly associated in only Olink-based but not SomaScan-based results. The blue color indicates CHIP variable-protein pairs significantly associated in only SomaScan-based but not Olink-based results. A total of 83 CHIP variable-protein pairs were significantly associated in both UKB and meta-analysis at P=0.05 threshold, and 16 of them were significant at FDR=0.05 level. ARIC: Atherosclerosis Risk in Communities; CHIP, clonal hematopoiesis of indeterminate potential; CHS: Cardiovascular Heart Study; EA: European ancestry; MESA: Multi-Ethnic Study of Atherosclerosis; UKB: UK Biobank.

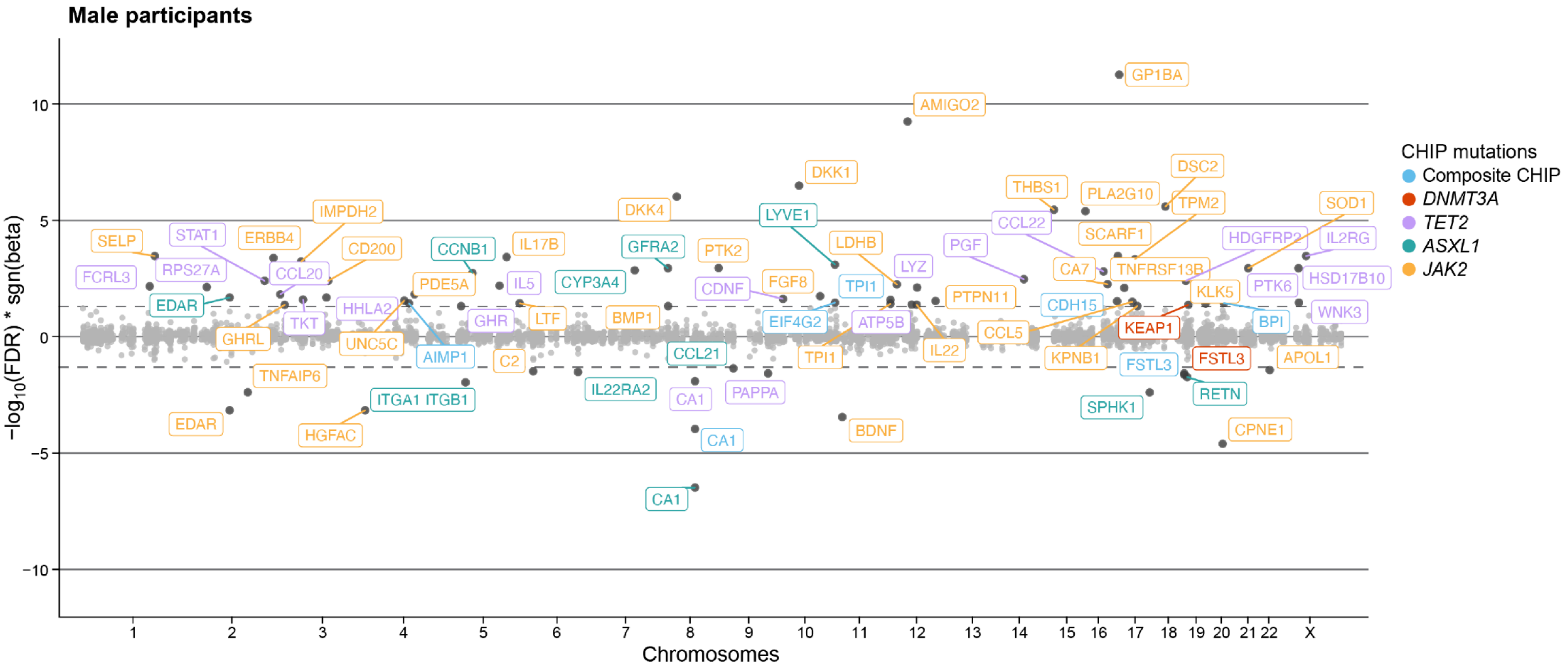


**Supplemental Figure 8. Meta-analyzed associations between CHIP mutations and circulating proteome among male participants in TOPMed cohorts** (N=5,616). Proteins that are associated at FDR=0.05 level (for 5,740 testings) are labeled with the corresponding SomaScan targets and colored in blue, red, green, orange, and purple, indicating significant associations with composite CHIP, DNMT3A, TET2, ASXL1, and JAK2, respectively. JAK2 analyses were only conducted in cohorts with greater than 5 participants with JAK2 mutations (i.e., CHS and ARIC EA). Associations were assessed through linear regression models adjusting for age at sequencing, race, batch (if applicable), type 2 diabetes status, smoker status, first ten principal components of genetic ancestry, and PEER factors (the number of PEER factors varies by cohorts based on the sizes of study populations: 50 for JHS, MESA, and CHS; 70 for ARIC AA; 120 for ARIC EA). AA: African ancestry; ARIC: Atherosclerosis Risk in Communities; CHIP, clonal hematopoiesis of indeterminate potential; CHS: Cardiovascular Heart Study; EA: European ancestry; FDR: false discovery rate; JHS: Jackson Heart Study; MESA: Multi-Ethnic Study of Atherosclerosis; PEER: probabilistic estimation of expression residuals.

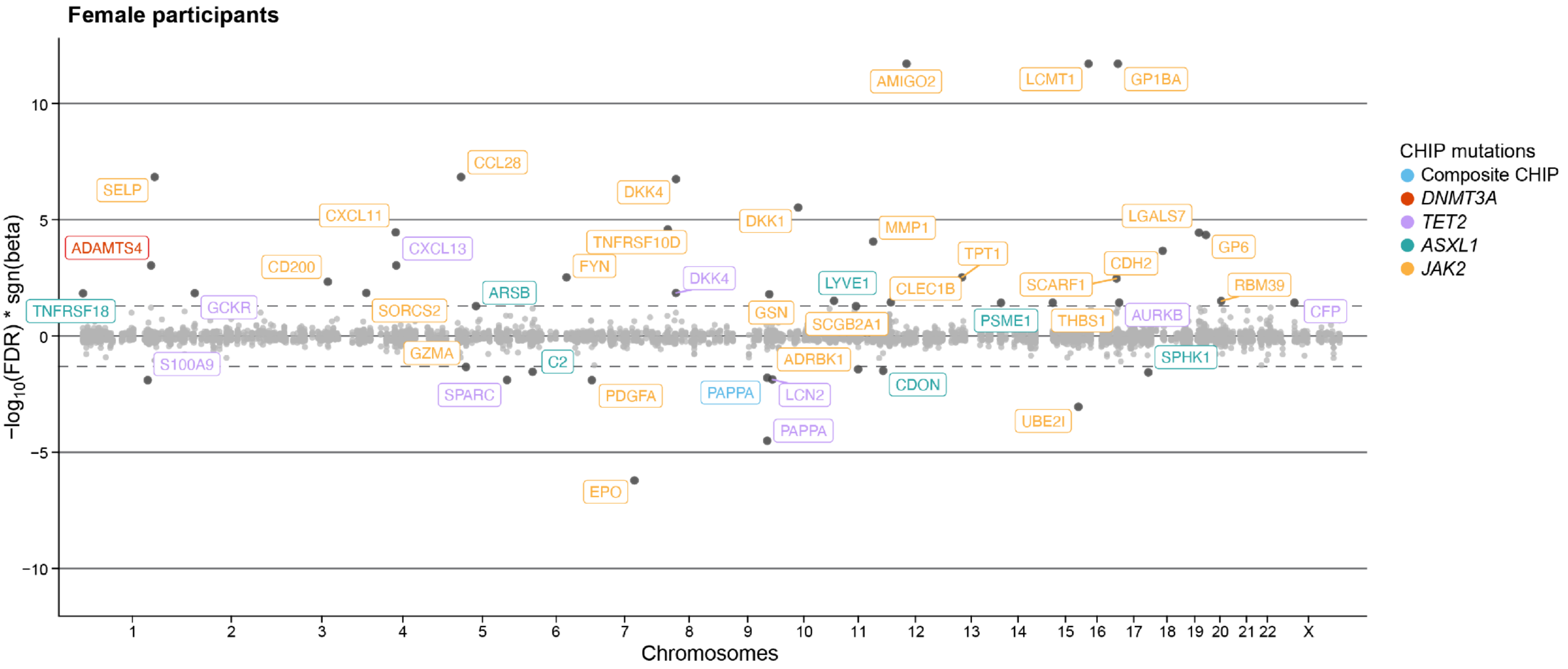


**Supplemental Figure 9. Meta-analyzed associations between CHIP mutations and circulating proteome among female participants in TOPMed cohorts** (N=7,295). Proteins that are associated at FDR=0.05 level (for 5,740 testings) are labeled with the corresponding SomaScan targets and colored in blue, red, green, orange, and purple, indicating significant associations with composite CHIP, DNMT3A, TET2, ASXL1, and JAK2, respectively. JAK2 analyses were only conducted in cohorts with greater than 5 participants with JAK2 mutations (i.e., CHS and ARIC EA). Associations were assessed through linear regression models adjusting for age at sequencing, race, batch (if applicable), type 2 diabetes status, smoker status, first ten principal components of genetic ancestry, and PEER factors (the number of PEER factors varies by cohorts based on the sizes of study populations: 50 for JHS, MESA, and CHS; 70 for ARIC AA; 120 for ARIC EA). AA: African ancestry; ARIC: Atherosclerosis Risk in Communities; CHIP, clonal hematopoiesis of indeterminate potential; CHS: Cardiovascular Heart Study; EA: European ancestry; FDR: false discovery rate; JHS: Jackson Heart Study; MESA: Multi-Ethnic Study of Atherosclerosis; PEER: probabilistic estimation of expression residuals.

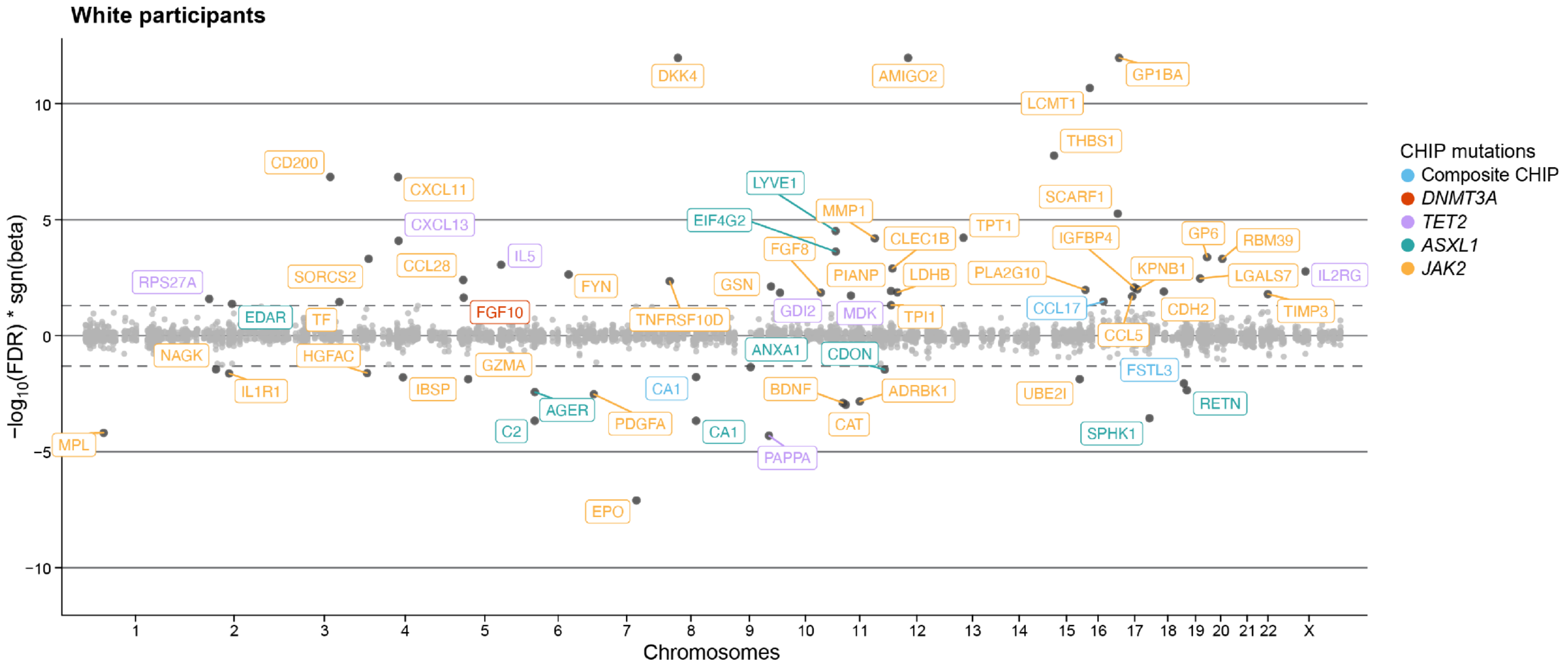


**Supplemental Figure 10. Meta-analyzed associations between CHIP mutations and circulating proteome among White participants in TOPMed cohorts (N=8,076).** Proteins that are associated at FDR=0.05 level (for 5,740 testings) are labeled with the corresponding SomaScan targets and colored in blue, red, green, orange, and purple, indicating significant associations with composite CHIP, DNMT3A, TET2, ASXL1, and JAK2, respectively. JAK2 analyses were only conducted in cohorts with greater than 5 participants with JAK2 mutations (i.e., CHS and ARIC EA). Associations were assessed through linear regression models adjusting for age at sequencing, sex, batch (if applicable), type 2 diabetes status, smoker status, first ten principal components of genetic ancestry, and PEER factors (the number of PEER factors varies by cohorts based on the sizes of study populations: 50 for JHS, MESA, and CHS; 120 for ARIC EA). ARIC: Atherosclerosis Risk in Communities; CHIP: Clonal hematopoiesis of indeterminate potential; CHS: Cardiovascular Heart Study; EA: European ancestry; FDR: False discovery rate; JHS: Jackson Heart Study; MESA: Multi-Ethnic Study of Atherosclerosis; PEER: Probabilistic estimation of expression residuals.

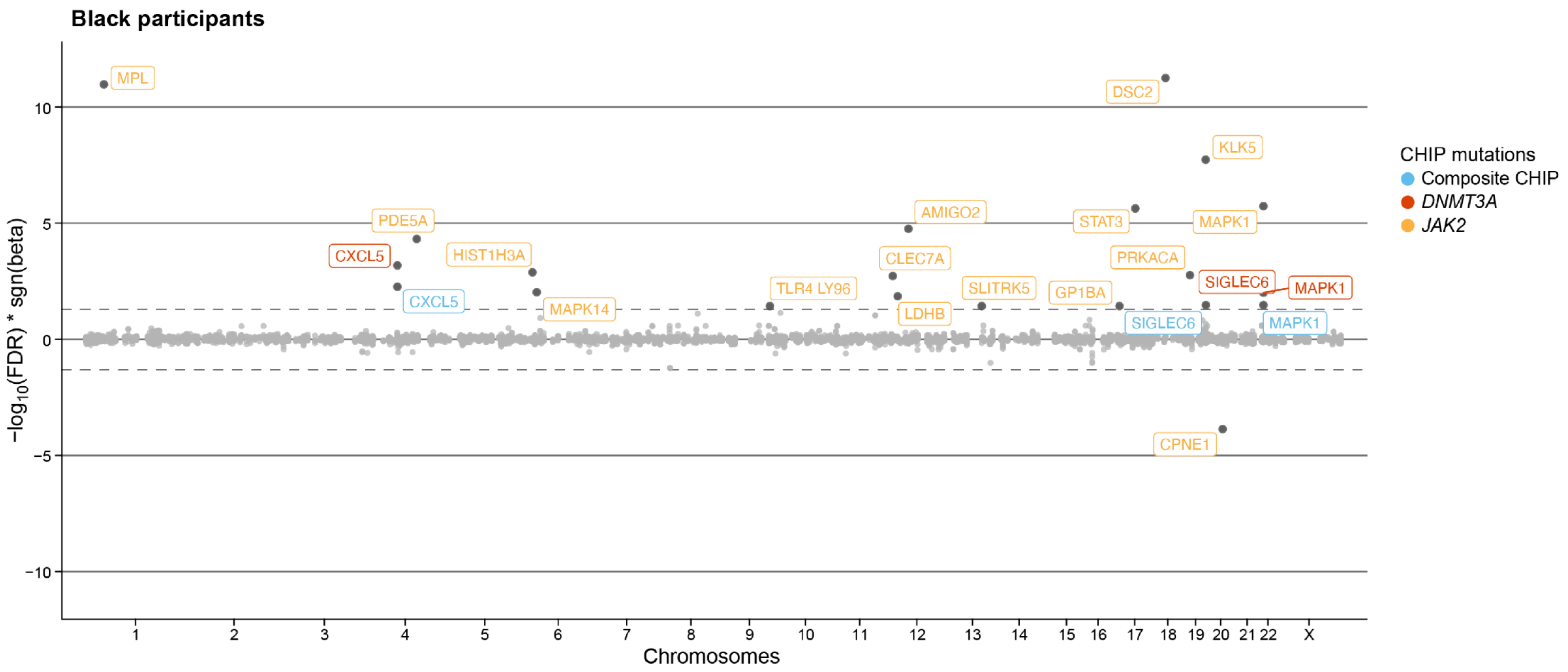


**Supplemental Figure 11. Meta-analyzed associations between CHIP mutations and circulating proteome among Black participants in TOPMed cohorts** (N=4,452). Proteins that are associated at FDR=0.05 level (for 5,740 testings) are labeled with the corresponding SomaScan targets and colored in blue, red, green, orange, and purple, indicating significant associations with composite CHIP, DNMT3A, TET2, ASXL1, and JAK2, respectively. JAK2 analyses were only conducted in cohorts with greater than 5 participants with JAK2 mutations (i.e., CHS and ARIC EA). Associations were assessed through linear regression models adjusting for age at sequencing, sex, batch (if applicable), type 2 diabetes status, smoker status, first ten principal components of genetic ancestry, and PEER factors (the number of PEER factors varies by cohorts based on the sizes of study populations: 50 for JHS, MESA, and CHS; 70 for ARIC AA). AA: African ancestry; ARIC: Atherosclerosis Risk in Communities; CHIP: Clonal hematopoiesis of indeterminate potential; CHS: Cardiovascular Heart Study; FDR: False discovery rate; JHS: Jackson Heart Study; MESA: Multi-Ethnic Study of Atherosclerosis; PEER: Probabilistic estimation of expression residuals.

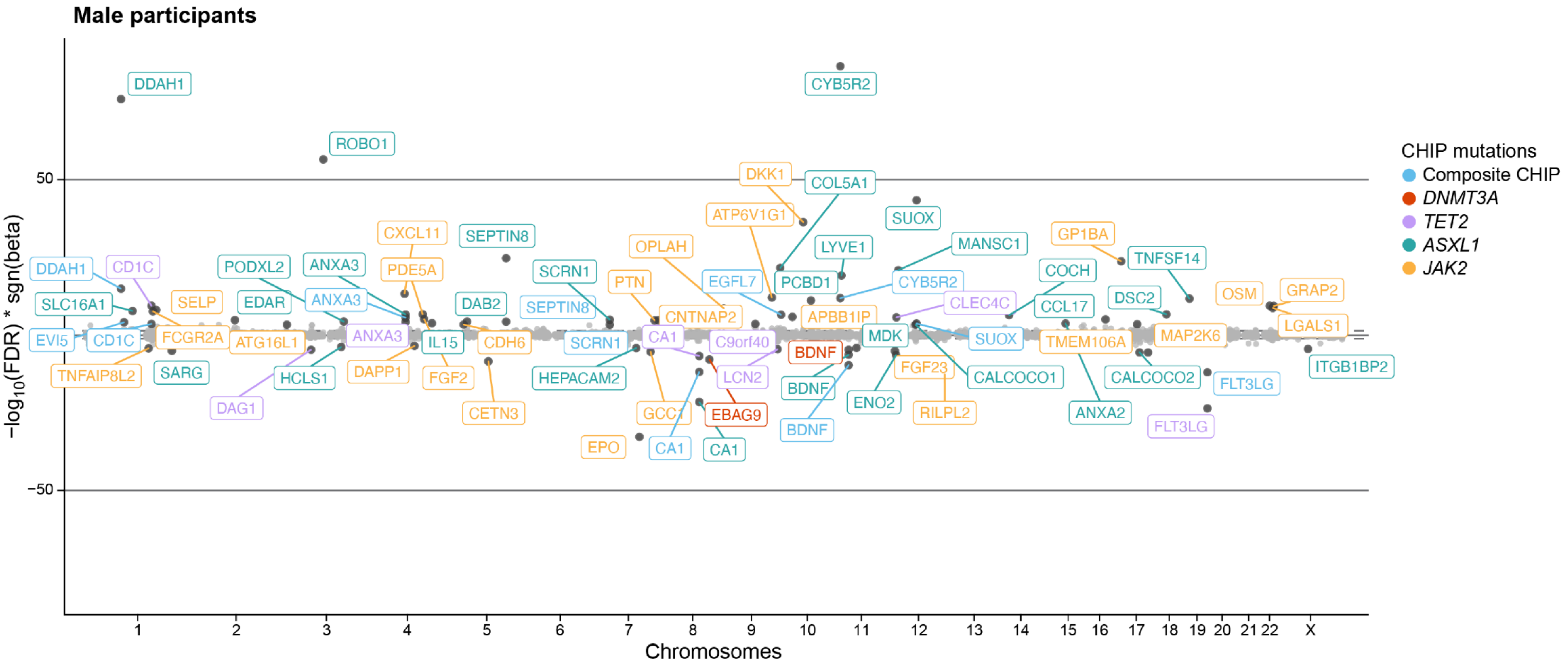


**Supplemental Figure 12. Associations between CHIP mutations and circulating proteome among male participants in UK Biobank (N=18,831).** Proteins that are associated at FDR=0.005 level (for 14,585 testings) are labeled with the corresponding Olink targets and colored in blue, red, purple, green, and orange, indicating significant associations with composite CHIP, *DNMT3A*, *TET2*, *ASXL1*, and *JAK2*, respectively. Associations were assessed through linear regression models adjusting for age at sequencing, self-reported British White ancestry (if applicable), type 2 diabetes status, current smoker status, first ten principal components of genetic ancestry, and 150 PEER factors. CHIP: Clonal hematopoiesis of indeterminate potential; FDR: False discovery rate; PEER: Probabilistic estimation of expression residuals.

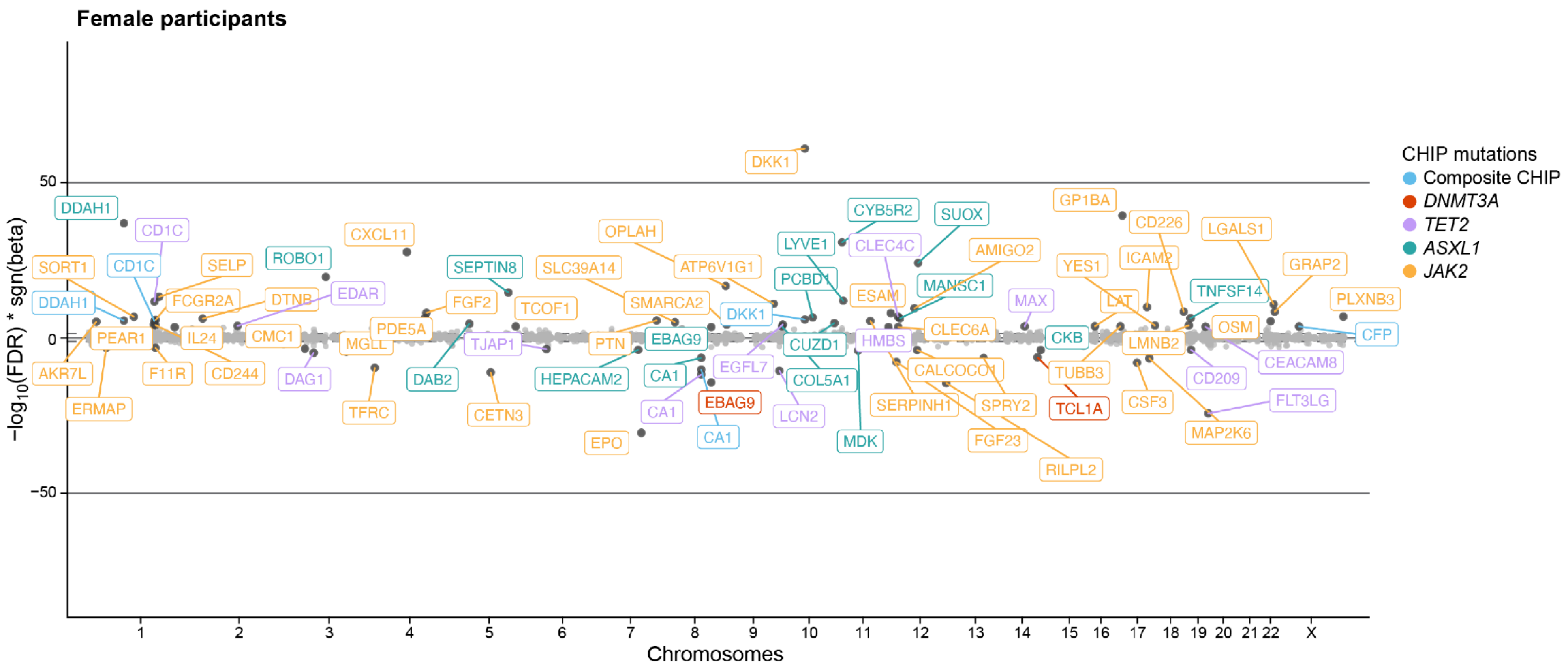


**Supplemental Figure 13. Associations between CHIP mutations and circulating proteome among female participants in UK Biobank (N=22,191).** Proteins that are associated at FDR=0.005 level (for 14,585 testings) are labeled with the corresponding Olink targets and colored in blue, red, purple, green, and orange, indicating significant associations with composite CHIP, *DNMT3A*, *TET2*, *ASXL1*, and *JAK2*, respectively. Associations were assessed through linear regression models adjusting for age at sequencing, self-reported British White ancestry (if applicable), type 2 diabetes status, current smoker status, first ten principal components of genetic ancestry, and 150 PEER factors. CHIP: Clonal hematopoiesis of indeterminate potential; FDR: False discovery rate; PEER: Probabilistic estimation of expression residuals.

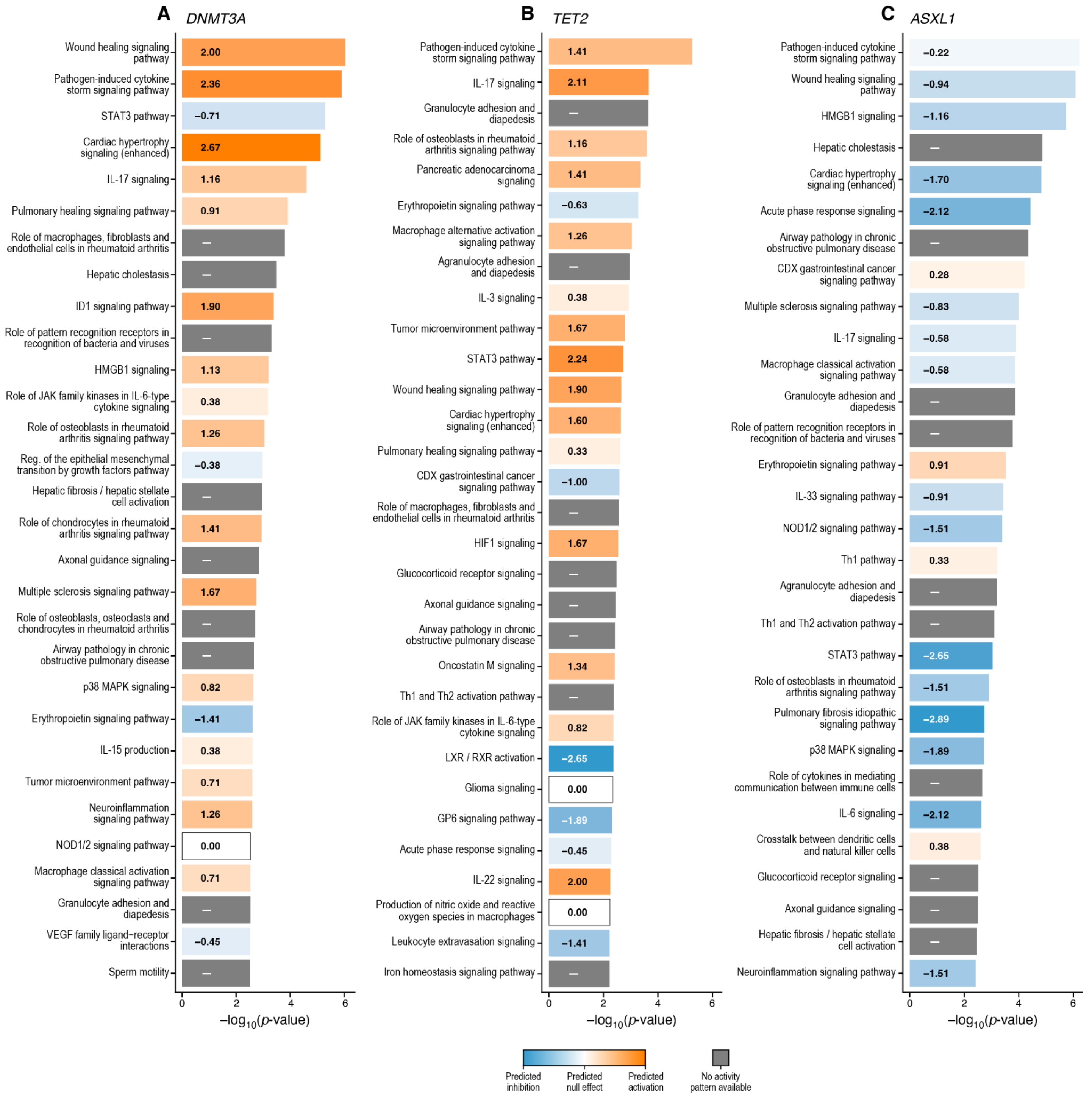

A DNMT3A
Wound healing signaling pathway 2.00
Pathogen-induced cytokine storm signaling pathway 2.36
STAT3 pathway −0.71
Cardiac hypertrophy signaling (enhanced) 2.67
IL-17 signaling 1.16
Pulmonary healing signaling pathway 0.91
Role of macrophages, fibroblasts and endothelial cells in rheumatoid arthritis –
Hepatic cholestasis –
ID1 signaling pathway 1.90
Role of pattern recognition receptors in recognition of bacteria and viruses –
HMGB1 signaling 1.13
Role of JAK family kinases in IL-6-type cytokine signaling 0.38
Role of osteoblasts in rheumatoid arthritis signaling pathway 1.26
Reg. of the epithelial mesenchymal transition by growth factors pathway −0.38
Hepatic fibrosis / hepatic stellate cell activation –
Role of chondrocytes in rheumatoid arthritis signaling pathway 1.41
Axonal guidance signaling –
Multiple sclerosis signaling pathway 1.67
Role of osteoblasts, osteoclasts and chondrocytes in rheumatoid arthritis –
Airway pathology in chronic obstructive pulmonary disease –
p38 MAPK signaling 0.82
Erythropoietin signaling pathway −1.41
IL-15 production 0.38
Tumor microenvironment pathway 0.71
Neuroinflammation signaling pathway 1.26
NOD1/2 signaling pathway 0.00
Macrophage classical activation signaling pathway 0.71
Granulocyte adhesion and diapedesis –
VEGF family ligand−receptor interactions −0.45
Sperm motility –
0 2 4 6
−log10(p-value)
B TET2
Pathogen-induced cytokine storm signaling pathway 1.41
IL-17 signaling 2.11
Granulocyte adhesion and diapedesis –
Role of osteoblasts in rheumatoid arthritis signaling pathway 1.16
Pancreatic adenocarcinoma signaling 1.41
Erythropoietin signaling pathway −0.63
Macrophage alternative activation signaling pathway 1.26
Agranulocyte adhesion and diapedesis –
IL-3 signaling 0.38
Tumor microenvironment pathway 1.67
STAT3 pathway 2.24
Wound healing signaling pathway 1.90
Cardiac hypertrophy signaling (enhanced) 1.60
Pulmonary healing signaling pathway 0.33
CDX gastrointestinal cancer signaling pathway −1.00
Role of macrophages, fibroblasts and endothelial cells in rheumatoid arthritis –
HIF1 signaling 1.67
Glucocorticoid receptor signaling –
Axonal guidance signaling –
Airway pathology in chronic obstructive pulmonary disease –
Oncostatin M signaling 1.34
Th1 and Th2 activation pathway –
Role of JAK family kinases in IL-6-type cytokine signaling 0.82
LXR / RXR activation −2.65
Glioma signaling 0.00
GP6 signaling pathway −1.89
Acute phase response signaling −0.45
IL-22 signaling 2.00
Production of nitric oxide and reactive oxygen species in macrophages 0.00
Leukocyte extravasation signaling −1.41
Iron homeostasis signaling pathway –
0 2 4 6
−log10(p-value)
C ASXL1
Pathogen-induced cytokine storm signaling pathway −0.22
Wound healing signaling pathway −0.94
HMGB1 signaling −1.16
Hepatic cholestasis –
Cardiac hypertrophy signaling (enhanced) −1.70
Acute phase response signaling −2.12
Airway pathology in chronic obstructive pulmonary disease –
CDX gastrointestinal cancer signaling pathway 0.28
Multiple sclerosis signaling pathway −0.83
IL-17 signaling −0.58
Macrophage classical activation signaling pathway −0.58
Granulocyte adhesion and diapedesis –
Role of pattern recognition receptors in recognition of bacteria and viruses –
Erythropoietin signaling pathway 0.91
IL-33 signaling pathway −0.91
NOD1/2 signaling pathway −1.51
Th1 pathway 0.33
Agranulocyte adhesion and diapedesis –
Th1 and Th2 activation pathway –
STAT3 pathway −2.65
Role of osteoblasts in rheumatoid arthritis signaling pathway −1.51
Pulmonary fibrosis idiopathic signaling pathway −2.89
p38 MAPK signaling −1.89
Role of cytokines in mediating communication between immune cells –
IL-6 signaling −2.12
Crosstalk between dendritic cells and natural killer cells 0.38
Glucocorticoid receptor signaling –
Axonal guidance signaling –
Hepatic fibrosis / hepatic stellate cell activation –
Neuroinflammation signaling pathway −1.51
0 2 4 6
−log10(p-value)
Predicted inhibition
Predicted null effect
Predicted activation
No activity pattern available

**Supplemental Figure 14. Significantly enriched pathways identified among proteins associated with CHIP driver genes.** Significantly enriched pathways corresponding to CHIP-associated proteins were derived based on known genetic and molecular relationships using IPA. The input was the Z-scores of the associations between major CHIP driver genes, i.e., *DNMT3A*, *TET2*, and *ASXL1*, and proteins that were significant at the P=0.05 level. The listed pathways are within the top 30 most significantly enriched pathways by input proteins based on IPA analysis (P<0.05). The orange indicates predicted activation, the blue indicates predicted inhibition, and the grey indicates no activity pattern available. The darker the orange and blue colors, the stronger the modulation effects. A. Canonical pathways implicated among proteins associated with *DNMT3A*. B. Canonical pathways implicated among proteins associated with *TET2*. C. Canonical pathways implicated among proteins associated with *ASXL1*. CHIP: Clonal hematopoiesis of indeterminate potential; IPA: Ingenuity Pathway Analysis

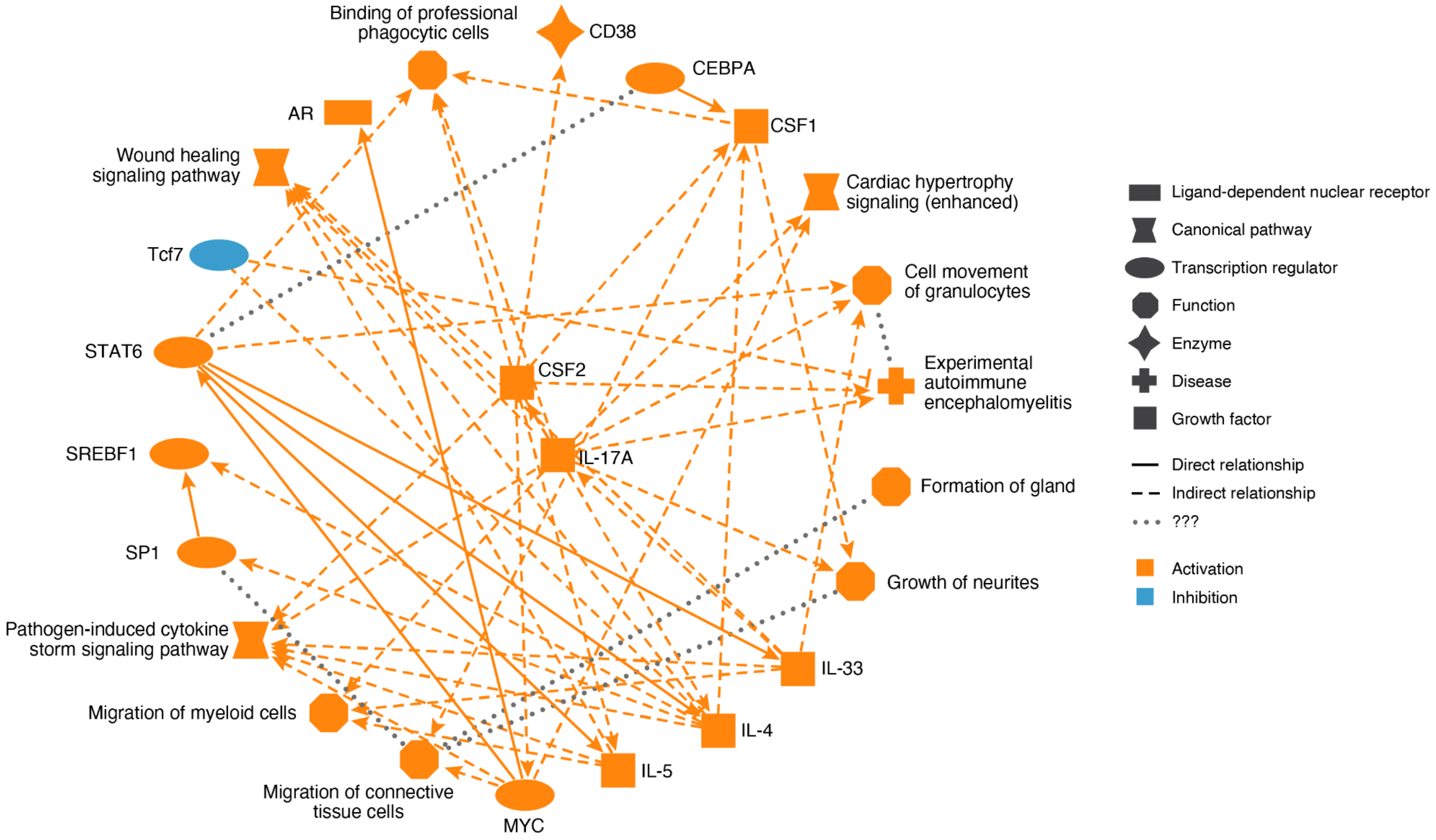


**Supplemental Figure 15. Summary of major biological themes for proteins associated with *DNMT3A*.** Biological themes of *DNMT3A*-associated proteins were derived based on known genetic and molecular relationships using IPA. The input was the Z-scores of the associations between *DNMT3A* and proteins that were significant at the P=0.05 level. The orange color indicates activation, and the blue color indicates inhibition. IPA: Ingenuity Pathway Analysis

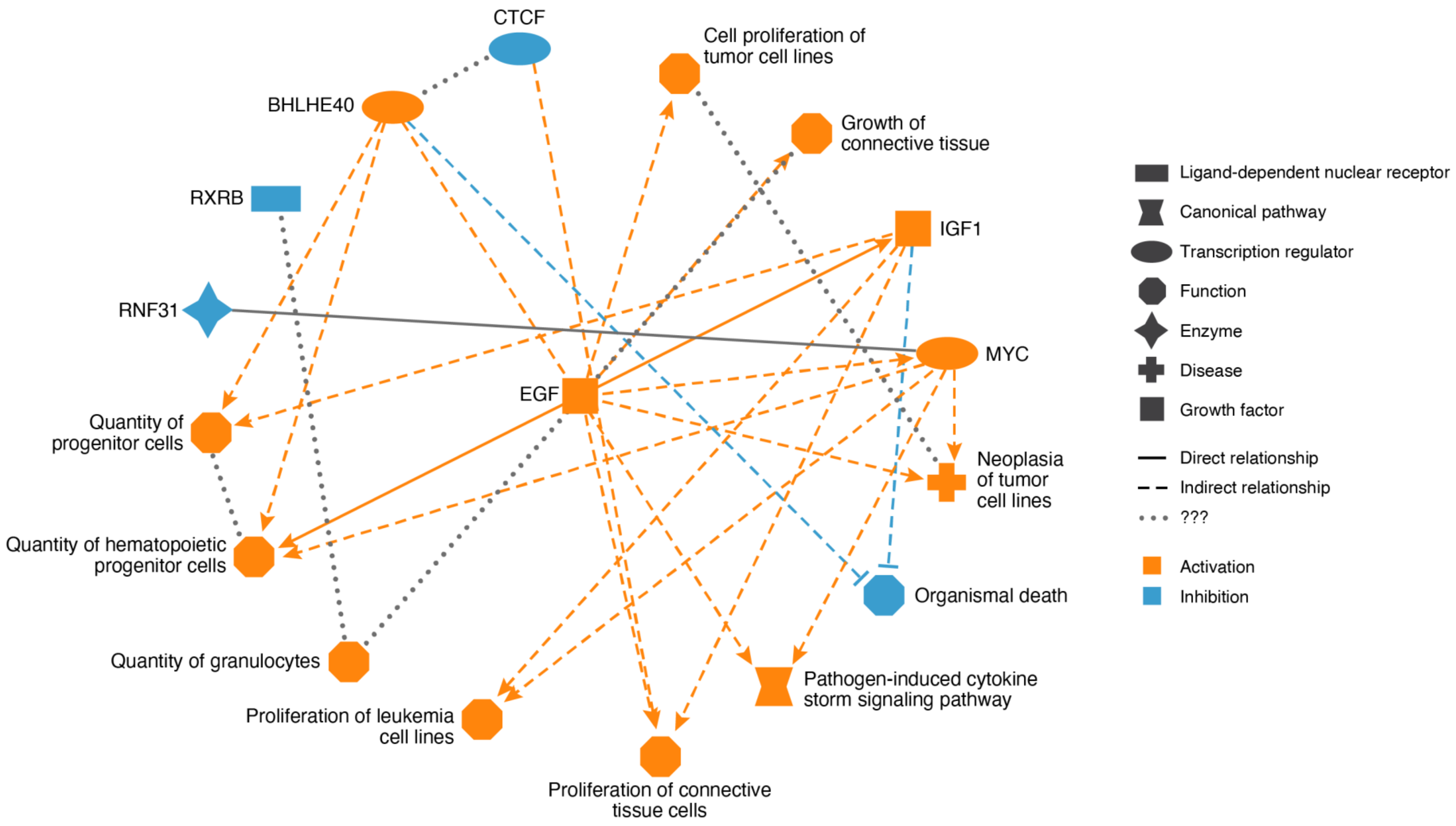


**Supplemental Figure 16. Summary of major biological themes for proteins associated with *TET2*.** Biological themes of *TET2*-associated proteins were derived based on known genetic and molecular relationships using IPA. The input was the Z-scores of the associations between *TET2* and proteins that were significant at the P=0.05 level. The orange color indicates activation, and the blue color indicates inhibition. IPA: Ingenuity Pathway Analysis.

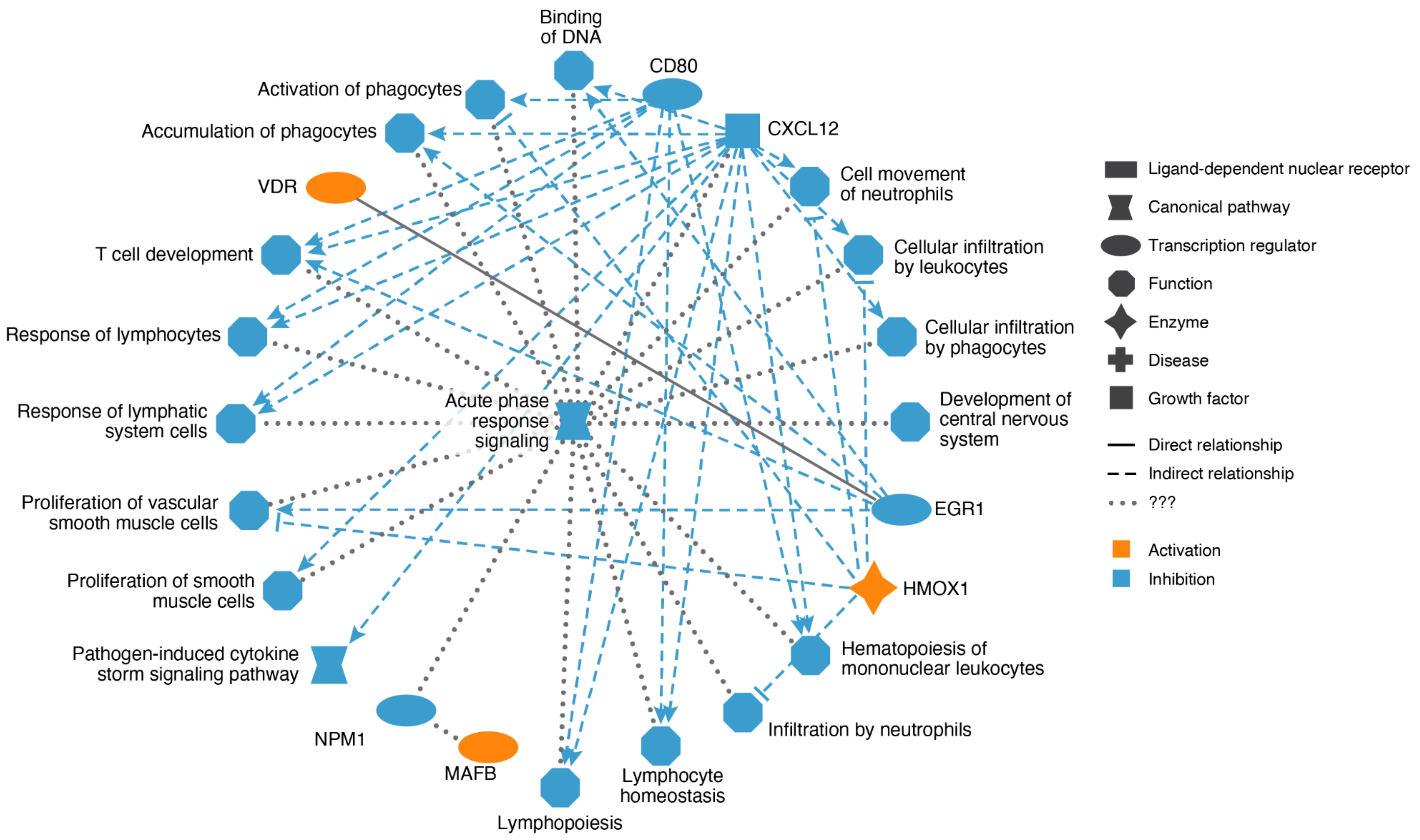


**Supplemental Figure 17. Summary of major biological themes for proteins associated with *ASXL1*.** Biological themes of *ASXL1*-associated proteins were derived based on known genetic and molecular relationships using IPA. The input was the Z-scores of the associations between *ASXL1* and proteins that were significant at the P=0.05 level. The orange color indicates activation, and the blue color indicates inhibition. IPA: Ingenuity Pathway Analysis.

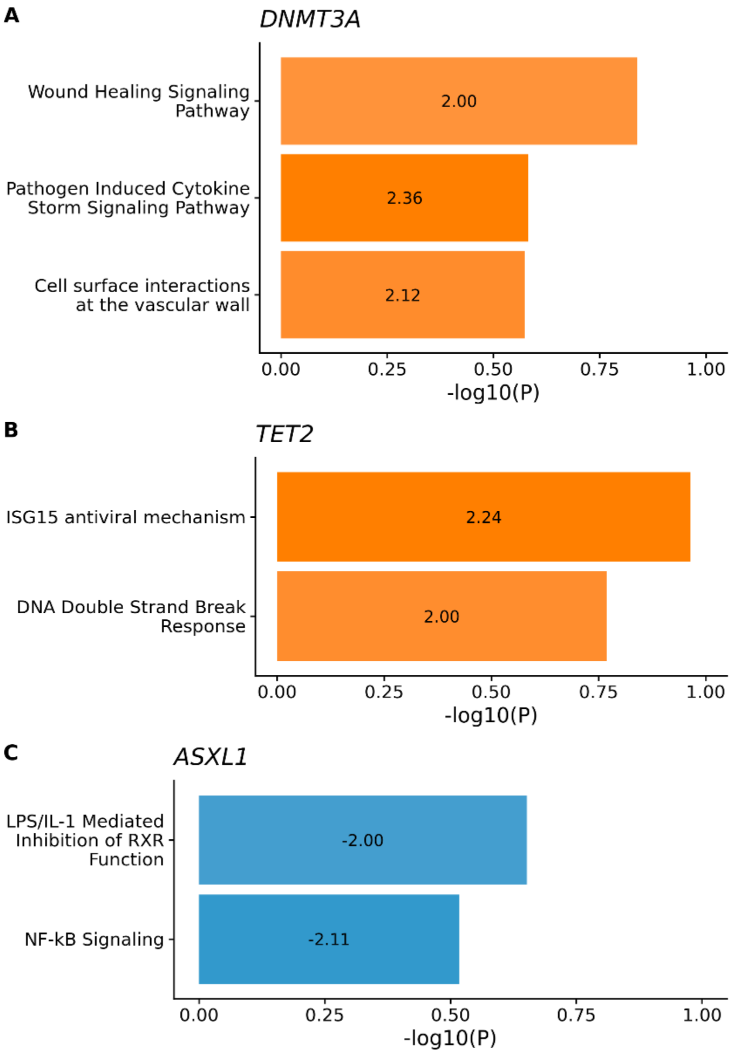


**Supplemental Figure 18. Significantly enriched and modulated pathways based on total quantified proteins were identified among proteins associated with CHIP driver genes.** Significantly enriched and modulated pathways corresponding to CHIP-associated proteins were derived based on known genetic and molecular relationships using IPA. We set the reference set as the total quantified proteins from our data. The input was the Z-scores of the associations between major CHIP driver genes, i.e., *DNMT3A*, *TET2*, and *ASXL1*, and proteins that were significant at the P=0.05 level. The listed pathways fulfill two criteria: (1) within the top 30 most significantly enriched pathways by input proteins based on IPA analysis (P<0.05) and (2) being significantly modulated, either inhibited or activated, based on IPA analysis (Z>1.96). The orange indicates predicted activation, and the blue indicates predicted inhibition. The darker the color, the stronger the modulation effect. A. Significantly modulated canonical pathways implicated among proteins associated with *DNMT3A*. B. Significantly modulated canonical pathways implicated among proteins associated with *TET2*. C. Significantly modulated canonical pathways implicated among proteins associated with *ASXL1*. CHIP: Clonal hematopoiesis of indeterminate potential; IPA: Ingenuity Pathway Analysis

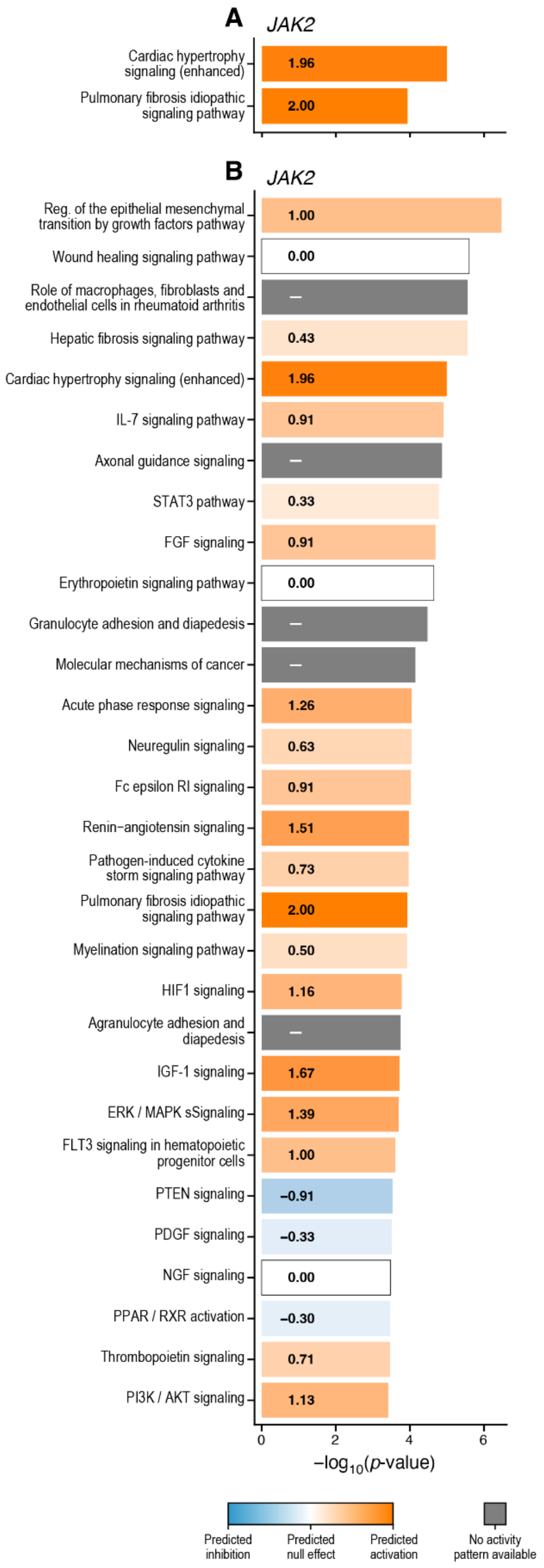


**Supplemental Figure 19. Significantly enriched and modulated pathways identified among proteins associated with *JAK2*.** Enriched and modulated pathways corresponding to *JAK2*-associated proteins were derived based on known genetic and molecular relationships using IPA. The input was the Z-scores of the associations between *JAK2* and proteins that were significant at the P=0.05 level. The listed pathways in A fulfill two criteria: (1) within the top 30 most significantly enriched pathways by input proteins based on IPA analysis ($P<0.05$) and (2) being significantly modulated, either inhibited or activated, based on IPA analysis ($Z>1.96$), and the listed pathways in B fulfills criteria (1). The orange indicates predicted activation, the blue indicates predicted inhibition, and the grey indicates no activity pattern available. The darker the color, the stronger the modulation effect. A. Significantly modulated canonical pathways implicated among proteins associated with *JAK2*. B. Canonical pathways implicated among proteins associated with *JAK2*. IPA: Ingenuity Pathway Analysis
