## Supplemental Text for "Human Plasma Proteomic Profile of Clonal Hematopoiesis"

**Protein Functions**

*TET2* CHIP-associated proteins discussed in the text:

- Pappalysin-1 (PAPPA; SomaScan target: PAPPA) is a metalloproteinase regulating insulin-like growth factor (IGF) by cleaving IGF-binding proteins. With modulating IGF availability, PAPPA plays crucial roles in fetal development[1-4], tissue remodeling, and regulation of various cellular processes[5-10].
- Secreted protein, acidic and rich in cysteine (SPARC; SomaScan target: ON), also called osteonectin or BM-40, is another matricellular glycoprotein that is involved in the development, remodeling, and tissue repair by modulating cell-cell and cell-matrix interactions[11-13].
- C-X-C motif chemokine 13 (CXCL13; SomaScan target: BLC) orchestrates B-cell trafficking to the site of inflammation and has been implicated in the development of several immune-related disorders[14-16].
- Neutrophil gelatinase-associated lipocalin (NGAL; SomaScan target: Lipocalin 2) is an innate immunity protein in response to pathologic states such as inflammation and infection[17]. It inhibits bacterial growth by binding siderophores with high affinity[18].
- Myeloperoxidase (SomaScan target: Myeloperoxidase) is a peroxidase enzyme involved in innate immunity and a marker for myeloid cells[19].

*ASXL1* CHIP-associated proteins discussed in the text:

- Carbonic anhydrase 1 (CA1; SomaScan target: Carbonic anhydrase I) is a member of the carbonic anhydrases family that mediates the reversible hydration of $CO_2$ to $H^+$ and $HCO_3^-$ and plays an essential role in maintaining acid-base balance[20]. CA1 is one of the 12

CAs that are present in humans and is localized in the cytosol of erythrocytes[21], neutrophils[22], cardiac tissues, gastrointestinal tract[23], and other organs and tissues.

- Resistin (SomaScan target: Resistin) is a hormone secreted from adipose tissue that resists insulin action and impairs glucose homeostasis. It has been implicated as the connecting link between visceral obesity and diabetes[24].
- Sphingosine kinase 1 (SPHK1; SomaScan target: Sphingosine kinase 1) is a cell-signaling enzyme localized mainly in the cytosol that mediates the conversion of ceramide to sphingosine-1-phosphate (S1P), which has been shown to induce tumor cell proliferation, resistance to chemotherapy, and metastasis[25].
- Lymphatic vessel endothelial hyaluronic acid receptor 1 (LYVE1; SomaScan target: LYVE1) is the main hyaluronan receptor in lymphatic vessel endothelium and has been used as a lymphatic-specific marker. It plays a key role in the trafficking of immunity-related cells, such as dendritic cells, macrophages, and leuckocytes, during inflammation, wound healing, and neoplasia[26,27].
- Cytochrome B5 reductase 2 (CYB5R2) is an enzyme involved in the electron transport chain that catalyzes the reduction of cytochrome b5. This process is essential for several metabolic pathways, including fatty acid desaturation and the metabolism of drugs and steroids, impacting cellular energy and overall metabolic homeostasis[28].
- Dimethylarginine dimethylaminohydrolase 1 (DDAH1) regulates nitric oxide production by metabolizing the endogenous nitric oxide synthase inhibitor asymmetric dimethylarginine (ADMA). This enzymatic activity is crucial for maintaining vascular tone and preventing endothelial dysfunction, thereby playing a vital role in cardiovascular health[29].

*JAK2* CHIP-associated proteins discussed in the text:

- P-selectin (SomaScan target: P-Selectin) is expressed on activated platelets and endothelial cells [30] and plays a role in the platelet-leukocyte interactions[31]. Its expression on platelets determines the size and stability of platelet aggregates[32]. It has been shown to contribute to pathological inflammation and thrombosis in many preclinical disease models, such as atherosclerosis and deep vein thrombosis[30].
- Platelet glycoprotein 1b alpha chain (GPIbα; SomaScan target: GP1BA) is a platelet surface membrane protein[33] that mediates the initial interaction with subendothelial von Willebrand factor (vWF)[34]. It has been implicated in atherothrombosis with genetic evidence supports the assertion that GPIbα influences atherothrombosis via increased platelet counts[35].
- Thrombospondin-1 (TSP1; SomaScan target: Thrombospondin-1) has been suggested as a counter receptor to GPIbα that supports initial platelet adhesion in the absence of vWF. It has also been reported that TSP1 modulates arterial thrombosis and contributes to arterial thrombosis in the presence of vWF[36]. TSP-1 is also an endogenous inhibitor of angiogenesis, inhibits endothelial function, stimulates apoptosis, and plays crucial functions in angiogenesis[37].

Discussed proteins in Black only analyses:

- Sialic acid-binding Ig-like lectin 6 (Siglec-6; SomaScan target: Siglec-6) is a member of the Siglec family of sialic acid-binding immunoglobulin-like lectins, which are expressed on various immune cells and selectively recognize different sialic acid-containing glycans[38]. The functions of Siglec-6 are not well understood, but it has been implicated in the modulation of immune tolerance.

- Mitogen-activated protein kinase 1 (MAK1; SomaScan target: MK01) is a protein kinase that plays a role in various cellular processes, including cell proliferation, differentiation, and survival. It is activated by phosphorylation and is involved in regulating gene expression in response to extracellular signals[39].

Examples of proteins showed alignment between human proteomic associations and mice differential expressions:

- Properdin (SomaScan target: Properdin) is a protein involved in the innate immune system as a key regulator of the alternative pathway (AP) of the complement system[40]. It is the only known positive regulator of AP and stabilizes the C3 convertase, increasing its half-life multiple times[41].
- Low-affinity immunoglobulin epsilon Fc receptor (FCER2; SomaScan target: CD23) is a membrane-bound protein predominantly expressed on B cells, monocytes, eosinophils, and other immune cells. It plays several critical roles in IgE-mediated immunity, including mediating the production of proinflammatory cytokines by monocytes[42], acting as a negative regulator for IgE and IgG antibody responses in B cells[43,44], etc.

References:

1. Lin, T.M., Galbert, S.P., Kiefer, D., Spellacy, W.N. & Gall, S. Characterization of four human pregnancy-associated plasma proteins. *Am J Obstet Gynecol* **118**, 223-236 (1974).
2. Wald, N., *et al.* First trimester concentrations of pregnancy associated plasma protein A and placental protein 14 in Down's syndrome. *Bmj* **305**, 28 (1992).
3. Kirkegaard, I., Uldbjerg, N. & Oxvig, C. Biology of pregnancy-associated plasma protein-A in relation to prenatal diagnostics: an overview. *Acta Obstet Gynecol Scand* **89**, 1118-1125 (2010).
4. Smith, G.C., *et al.* Early-pregnancy origins of low birth weight. *Nature* **417**, 916 (2002).
5. Boldt, H.B. & Conover, C.A. Overexpression of pregnancy-associated plasma protein-A in ovarian cancer cells promotes tumor growth in vivo. *Endocrinology* **152**, 1470-1478 (2011).
6. Dalgin, G.S., Holloway, D.T., Liou, L.S. & DeLisi, C. Identification and characterization of renal cell carcinoma gene markers. *Cancer Inform* **3**, 65-92 (2007).
7. Loddo, M., *et al.* Pregnancy-associated plasma protein A regulates mitosis and is epigenetically silenced in breast cancer. *J Pathol* **233**, 344-356 (2014).
8. Bulut, I., *et al.* Relationship between pregnancy-associated plasma protein-A and lung cancer. *Am J Med Sci* **337**, 241-244 (2009).
9. Nagarajan, N., *et al.* Whole-genome reconstruction and mutational signatures in gastric cancer. *Genome Biol* **13**, R115 (2012).
10. Huang, J., *et al.* Identification of pregnancy-associated plasma protein A as a migration-promoting gene in malignant pleural mesothelioma cells: a potential therapeutic target. *Oncotarget* **4**, 1172-1184 (2013).

11. Brekken, R.A. & Sage, E.H. SPARC, a matricellular protein: at the crossroads of cell-matrix communication. *Matrix Biol* **19**, 816-827 (2001).

12. Bradshaw, A.D. & Sage, E.H. SPARC, a matricellular protein that functions in cellular differentiation and tissue response to injury. *J Clin Invest* **107**, 1049-1054 (2001).

13. Tai, I.T. & Tang, M.J. SPARC in cancer biology: its role in cancer progression and potential for therapy. *Drug Resist Updat* **11**, 231-246 (2008).

14. Müller, G., Höpken, U.E. & Lipp, M. The impact of CCR7 and CXCR5 on lymphoid organ development and systemic immunity. *Immunol Rev* **195**, 117-135 (2003).

15. Meraouna, A., *et al.* The chemokine CXCL13 is a key molecule in autoimmune myasthenia gravis. *Blood* **108**, 432-440 (2006).

16. Lee, H.T., *et al.* Serum BLC/CXCL13 concentrations and renal expression of CXCL13/CXCR5 in patients with systemic lupus erythematosus and lupus nephritis. *J Rheumatol* **37**, 45-52 (2010).

17. Kjeldsen, L., Cowland, J.B. & Borregaard, N. Human neutrophil gelatinase-associated lipocalin and homologous proteins in rat and mouse. *Biochim Biophys Acta* **1482**, 272-283 (2000).

18. Yang, J., *et al.* An iron delivery pathway mediated by a lipocalin. *Mol Cell* **10**, 1045-1056 (2002).

19. Klebanoff, S.J. Myeloperoxidase: friend and foe. *J Leukoc Biol* **77**, 598-625 (2005).

20. Lindskog, S. Structure and mechanism of carbonic anhydrase. *Pharmacol Ther* **74**, 1-20 (1997).

21. Konialis, C.P., Barlow, J.H. & Butterworth, P.H. Cloned cDNA for rabbit erythrocyte carbonic anhydrase I: A novel erythrocyte-specific probe to study development in erythroid tissues. *Proc Natl Acad Sci U S A* **82**, 663-667 (1985).

22. Campbell, A.R., Andress, D.L. & Swenson, E.R. Identification and characterization of human neutrophil carbonic anhydrase. *J Leukoc Biol* **55**, 343-348 (1994).

23. Parkkila, S., Parkkila, A.K., Juvonen, T. & Rajaniemi, H. Distribution of the carbonic anhydrase isoenzymes I, II, and VI in the human alimentary tract. *Gut* **35**, 646-650 (1994).

24. Steppan, C.M., *et al.* The hormone resistin links obesity to diabetes. *Nature* **409**, 307-312 (2001).

25. Ogretmen, B. Sphingolipid metabolism in cancer signalling and therapy. *Nat Rev Cancer* **18**, 33-50 (2018).

26. Jackson, D.G. Hyaluronan in the lymphatics: The key role of the hyaluronan receptor LYVE-1 in leucocyte trafficking. *Matrix Biol* **78-79**, 219-235 (2019).

27. Lee, J.Y. & Spicer, A.P. Hyaluronan: a multifunctional, megaDalton, stealth molecule. *Curr Opin Cell Biol* **12**, 581-586 (2000).

28. Hall, R., Yuan, S., Wood, K., Katona, M. & Straub, A.C. Cytochrome b5 reductases: Redox regulators of cell homeostasis. *J Biol Chem* **298**, 102654 (2022).

29. Murphy, R.B., Tommasi, S., Lewis, B.C. & Mangoni, A.A. Inhibitors of the Hydrolytic Enzyme Dimethylarginine Dimethylaminohydrolase (DDAH): Discovery, Synthesis and Development. *Molecules* **21**(2016).

30. McEver, R.P. Selectins: initiators of leucocyte adhesion and signalling at the vascular wall. *Cardiovasc Res* **107**, 331-339 (2015).

31. Totani, L. & Evangelista, V. Platelet-leukocyte interactions in cardiovascular disease and beyond. *Arterioscler Thromb Vasc Biol* **30**, 2357-2361 (2010).

32. Merten, M. & Thiagarajan, P. P-selectin expression on platelets determines size and stability of platelet aggregates. *Circulation* **102**, 1931-1936 (2000).

33. Luo, S.Z., *et al.* Glycoprotein Ibalpha forms disulfide bonds with 2 glycoprotein Ibbeta subunits in the resting platelet. *Blood* **109**, 603-609 (2007).

34. Bergmeier, W., *et al.* The role of platelet adhesion receptor GPIbalpha far exceeds that of its main ligand, von Willebrand factor, in arterial thrombosis. *Proc Natl Acad Sci U S A* **103**, 16900-16905 (2006).

35. Sun, B.B., *et al.* Genomic atlas of the human plasma proteome. *Nature* **558**, 73-79 (2018).

36. Prakash, P., Kulkarni, P.P. & Chauhan, A.K. Thrombospondin 1 requires von Willebrand factor to modulate arterial thrombosis in mice. *Blood* **125**, 399-406 (2015).

37. Zhao, C., Isenberg, J.S. & Popel, A.S. Human expression patterns: qualitative and quantitative analysis of thrombospondin-1 under physiological and pathological conditions. *J Cell Mol Med* **22**, 2086-2097 (2018).

38. Crocker, P.R., Paulson, J.C. & Varki, A. Siglecs and their roles in the immune system. *Nat Rev Immunol* **7**, 255-266 (2007).

39. Cargnello, M. & Roux, P.P. Activation and function of the MAPKs and their substrates, the MAPK-activated protein kinases. *Microbiol Mol Biol Rev* **75**, 50-83 (2011).

40. Walport, M.J. Complement. First of two parts. *N Engl J Med* **344**, 1058-1066 (2001).

41. Fearon, D.T. & Austen, K.F. Properdin: binding to C3b and stabilization of the C3b-dependent C3 convertase. *J Exp Med* **142**, 856-863 (1975).

42. Lecoanet-Henchoz, S., *et al.* CD23 regulates monocyte activation through a novel interaction with the adhesion molecules CD11b-CD18 and CD11c-CD18. *Immunity* **3**, 119-125 (1995).

43. Campbell, K.A., *et al.* Induction of B cell apoptosis by co-cross-linking CD23 and sIg involves aberrant regulation of c-myc and is inhibited by bcl-2. *Int Immunol* **9**, 1131-1140 (1997).

44. Payet, M.E., Woodward, E.C. & Conrad, D.H. Humoral response suppression observed with CD23 transgenics. *J Immunol* **163**, 217-223 (1999).
